# Designing Sub-20 nm Nanocarriers for Small Molecule Delivery: Interplay among Structural Geometry, Assembly Energetics, and Cargo Release Kinetics

**DOI:** 10.1101/2020.08.14.245340

**Authors:** Benson T. Jung, Marc Lim, Katherine Jung, Michael Li, He Dong, Nikhil Dube, Ting Xu

**Author notes:** These authors contributed equally to the manuscript (Benson T. Jung, Marc Lim). Department of Chemistry and Biochemistry, University of Texas at Arlington, 169 Science & Engineering Innovation & Research Building, Arlington, TX 76019, United States.

## Abstract

Biological constraints in diseased tissues have motivated the need for small nanocarriers (10-30 nm) to achieve sufficient vascular extravasation and pervasive tumor penetration. This particle size limit is only an order of magnitude larger than small molecules, such that cargo loading is better described by co-assembly processes rather than simple encapsulation. Understanding the structural, kinetic, and energetic contributions of carrier-cargo co-assembly is thus critical to achieve molecular-level control and predictable *in vivo* behavior. These interconnected set of properties were systematically examined using sub-20 nm self-assembled nanocarriers known as three-helix micelles (3HM). Both hydrophobicity and the “geometric packing parameter” dictate small molecule compatibility with 3HM’s alkyl tail core. Planar obelisk-like apomorphine and doxorubicin (DOX) molecules intercalated well within the 3HM core and near the core-shell interface, forming an integral component to the co-assembly, as corroborated by small angle X-ray and neutron-scattering structural studies. DOX promoted crystalline alkyl tail ordering, which significantly increased (+63%) the activation energy of 3HM subunit exchange. Subsequently, 3HM-DOX displayed slow-release kinetics (t_1/2_=40 h) at physiological temperatures, with ~50x greater cargo preference for the micelle core as described by two drug partitioning coefficients (micellar core/shell K_p1_ ~24, and shell/bulk solvent K_p2_ ~2). The geometric and energetic insights between nanocarrier and their small molecule cargos developed here will aid in broader efforts to deconvolute the interconnected properties of carrier-drug co-assemblies, and to understand nanomedicine behavior throughout all the physical and *in vivo* processes they are intended to encounter.

## 1. Introduction

Nanomedicine has long-promised enhanced therapeutic benefits such as increasing the accumulation of drugs in target tissues and decreasing systemic side effects. ^1–5^ Advancing physical design parameters is requisite toward successful clinical translation of nanomaterial therapies, which still suffer from diminishing success in the clinic (decreasing from 94% in Phase I to 14% in Phase III) ^6–10^ Disease pathology and nanomaterial research have motivated the need for small nanocarriers (10-30 nm) to conform to the physiological realities of drug delivery transport events, including vascular extravasation and pervasive tumor penetration. ^11–16^ At these length-scales, the relative impact of every molecular participant in a nanomedicine formulation becomes magnified. With nanocarrier feature sizes approaching those of the small molecule therapeutics (~1 nm) they are intended to deliver, it is necessary to consider cargo formulation less as a simple encapsulation, and more as a co-assembly process. ^17^ As volume decreases with r^3^, geometric considerations between cargo and amphiphiles become critical to particle structure (akin to the “critical packing parameter” ^18^), drug loading capacity, and kinetic stability (through examination of alkyl core phase behavior). ^19,20^ Yet, physicochemical studies at this level of molecular precision are scarce for nanocarrier co-assemblies in this size range.^11,12,21^ Understanding carrier-drug co-assembly will inform design rules for nanocarrier development, enabling rational optimization of carriers with predictable kinetic behavior under complex *in vivo* environments. ^10,21^ Achieving molecular-level understanding has benefited analogous fields such as supramolecular amphiphilic assembly, which evolved from simple 2D polymers to advanced dynamically-signaling biomaterials. ^22–27^ Knowing the precise location of Quantum Dots loaded in nanocarriers yielded improved stability, reduced toxicity and more reliable optical performance. ^28^ Shifting analytical approaches to quantitative analysis, is needed to deepened insights into nanocarrier stability and subunit interactions. ^29,30^ Especially when pursuing self-assembling systems, it becomes critical to consider 1) how drug cargo complexation affects small nanocarrier self-assembly, structure, and kinetic stability 2) how cargo selection, spatial distribution, and release are affected by restricted volume, and 3) how to quantify and control the thermodynamic energy landscape in multi-component systems. Advanced techniques such as small angle X-ray and neutron scattering allow detailed structural information to be gleaned. ^31,32^ They are valuable tools to further characterize the morphology, internal distribution, and interactions of these complex co-assemblies beyond characterization techniques commonly employed for nanoparticle analysis, such as transmission electron microscopy, dynamic light scattering, and zeta potential. ^31,33,34^

Here, fundamental insights into nanocarrier-cargo co-assembly are pursued using a sub-20 nm model particle known as 3-helix micelles (3HM). 3HM’s architectural components approach the dimensions of small molecule therapeutics **(Scheme 1 A**). 3HM has homogeneous particle size ~18 nm with high *in vitro* stability, and details of its assembly kinetic pathways, rate of monomer exchange, and internal structure has been quantitatively studied in-depth. ^4,5,35–39^ Additionally, 3HM has been shown to possess favorable *in vivo* biodistribution, pharmacokinetics, and accumulation in orthotopically xenografted brain tumors. ^4,5,35–37^ Energetic contributions from each of the comprising components (peptide,^37,40^ polymer, ^37,39^ and alkyl chains ^36^) have been systematically studied. Building upon this well-rounded background knowledge, 3HM becomes a great model system to investigate nanomaterial-drug co-assembly.

**Scheme 1:**
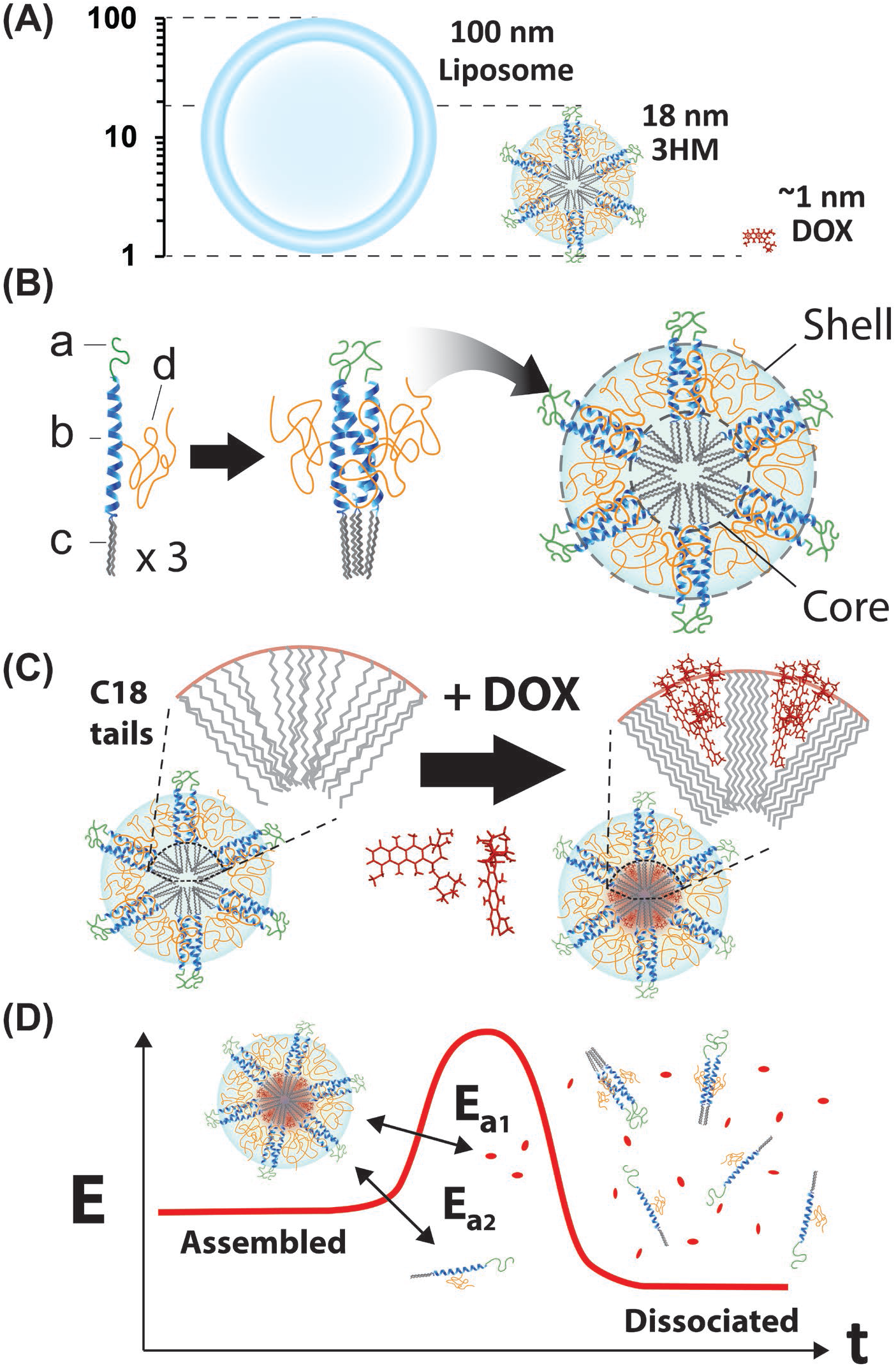
(A) Geometric scales comparing 100 nm liposome, 18 nm 3HM, and 1 nm small molecule such as DOX. (B) 3HM monomers are composed of peptide polymer and alkyl chains; a = polyethylene glycol (PEG) 750 Da, b = alpha-helix peptide, c = C18 alkyl tails, d = PEG 2000 Da. The peptides can form trimeric coiled-coils, which assemble into a micellar structure with hydrophobic core and hydrophilic shell. (C) In the 3HM core, C18 tails splayed out with a low degree of crystallinity gain well-ordered conformation upon DOX co-assembly, due to favorable drug geometry and hydrophobicity for intercalating between C18 tails in the core and at the core-shell interface. (D) The overall 3HM-DOX dissociation energetics encompasses both DOX and monomer desorption, leading to slow cargo release kinetics.

The present studies demonstrated that 3HM nanocarrier internal structure, cargo spatial distribution, and geometry, are tied to co-assembly energetics and eventual cargo release kinetics (**Scheme 1**). The “geometric packing parameter” of a small molecule is equally significant to its degree of hydrophobicity for optimizing 3HM cargo loading. The resultant co-assembly increased crystalline alkyl tail ordering as well as energetic barriers towards carrier-cargo disassembly, creating a slow-release system with high drug partitioning preference for the micellar core. Results presented here substantiate that at sub-20 nm length-scales, the concept of simple encapsulation needs to be revisited. The insights gained here should be applicable to other nanocarrier–drug combinations and provide much needed design guidelines. With parallel ongoing work to examine nanomaterial fate at the cellular and organism scales, knowledge here can be useful in endeavors to forecast nanomedicine behavior from benchtop formulation to *in vivo* excretion.

## 2. Experimental Methods

### 2.1: Synthesis of Amphiphilic Peptide−Polymer 3HM Conjugates

The synthetic and purification procedures of the 3HM amphiphile have been documented in detail previously. ^35,36^ Briefly, the 3-helix bundle forming peptide, 1CW (EVEALEKKVAALEC KVQALEKKVEALEHGW) was modified at the N-terminus with Fmoc-6-aminohexanoic acid (Ahx) as a linker followed by a Fmoc-Lys(Fmoc)-OH residue and at the C-terminus with additional residues Fmoc-Gly-Gly-Gly-Lys(Alloc)-OH. N-terminal modification allowed for conjugation of two alkyl tails leading to the amphiphilic molecule. Unless specified otherwise, stearic acid (C18) alkyl tails were primarily used in this study. Peptide C-terminal modification allowed for conjugation of carboxy-terminated PEG750, to provide a stealth layer on the surface-exposed C-terminus. The cysteine residue at position 14 facilitated conjugation of a maleimide-PEG2K to the amphiphile that provides entropic stabilization reported in detail previously. ^37^

Using a Prelude solid phase peptide synthesizer (Protein Technologies, AZ), the modified peptide (K(Fmoc)-Ahx-EVEALEKKVAALECKVQALEKKVEALEHGWGGGK(Alloc)) was produced via 9-fluorenylmethyl carbamate (Fmoc) chemistry. The N-terminal alkyl chains were conjugated through reaction of stearic acid with deprotected Fmoc-Lys(Fmoc)-OH. The C-terminal Fmoc-Lys(Alloc)-OH was selectively deprotected with five 30 min reactions of 0.2 molar equivalents of Pd(PPh_3_)_4_ used as a catalyst, with 24 equivalents of phenylsilane as an allyl acceptor in dichloromethane. The resulting free amino group was utilized for conjugating carboxy-terminated PEG750 using HBTU/DIPEA chemistry. Cleavage was carried out using a cocktail of 95:2.5:2.5 TFA/TIS/water for 3 h. Crude peptides were precipitated and washed three times in cold ether, isolated, and dried.

Maleimide-Cys conjugation was used to specifically conjugate PEG2K to the cysteine at position 14 in the middle of the peptide sequence. The conjugation reaction was carried out in 9:1 HEPES buffer (100 mM, pH=7.4):MeOH overnight under nitrogen with a reaction ratio of PEG to peptide at 2:1. The final peptide amphiphiles (MW= ~7200 g/mol) were isolated with centrifugal filters (Amicon, MW cutoff: 3000 g/mol). Spin filtration was performed at 7000 rpm for 40 min and the concentrate was washed with deionized water. This was repeated seven times, then the amphiphiles were lyophilized to a powder. For SANS studies, deuterated alkyl tails were used to provide contrast variation. Peptide amphiphile masses were confirmed with Matrix-Assisted Laser Desorption Ionization Time of Flight mass spectrometry.^35,38^

### 2.2: Cargo Co-Assembly

The thin film hydration method to co-assembly 3HM with DOX has been described previously. ^5^ As a system with inherent sensitivity to thermal history, ^38^ each step of the 3HM co-assembly process with DOX has the potential to change the structure and properties of the overall nanocarrier. Hence, in these studies the thin film rehydration process and intermediate stages leading toward the final product has been fixed (**Scheme 2**). Briefly, purified 3HM was dissolved in methanol to 20 mg/mL. 0.2 mg/mL DOX in methanol was added to 13 % w/w of 3HM. The co-dissolved 3HM and DOX was sonicated and left to dry slowly to a film under air flow in darkness at room temperature. Residual methanol was evaporated in a vacuum oven for 2h. Potassium phosphate buffer (25 mM KH_2_PO_4_, pH 7.4) was used to rehydrate the film to ~1 mg/mL. The micelle solution was annealed at 70 °C for 45 min and stirred at room temperature overnight to equilibrate the drug loaded micelles. Free DOX was removed with centrifugal filters (Amicon Ultra, MW cutoff: 3000 g/mol). Spin filtration was performed at 7500 rpm for 30 min and the retentate was washed with deionized water. This was repeated until DOX became virtually undetectable in the filtrate, whereby 480 nm UV-vis absorbance reached a constant low value of 0.02, usually after 7-10 times. This indicates complete removal of free DOX, with final drug loading at ~8-9 % w/w. The retained 3HM-DOX concentrate was lyophilized for storage. For further experiments, 3HM-DOX was dissolved in phosphate buffer (25 mM, pH 7.4) to the desired concentration, annealed at 70 °C for 45 min, and allowed to cool to room temperature before use.

**Scheme 2:**
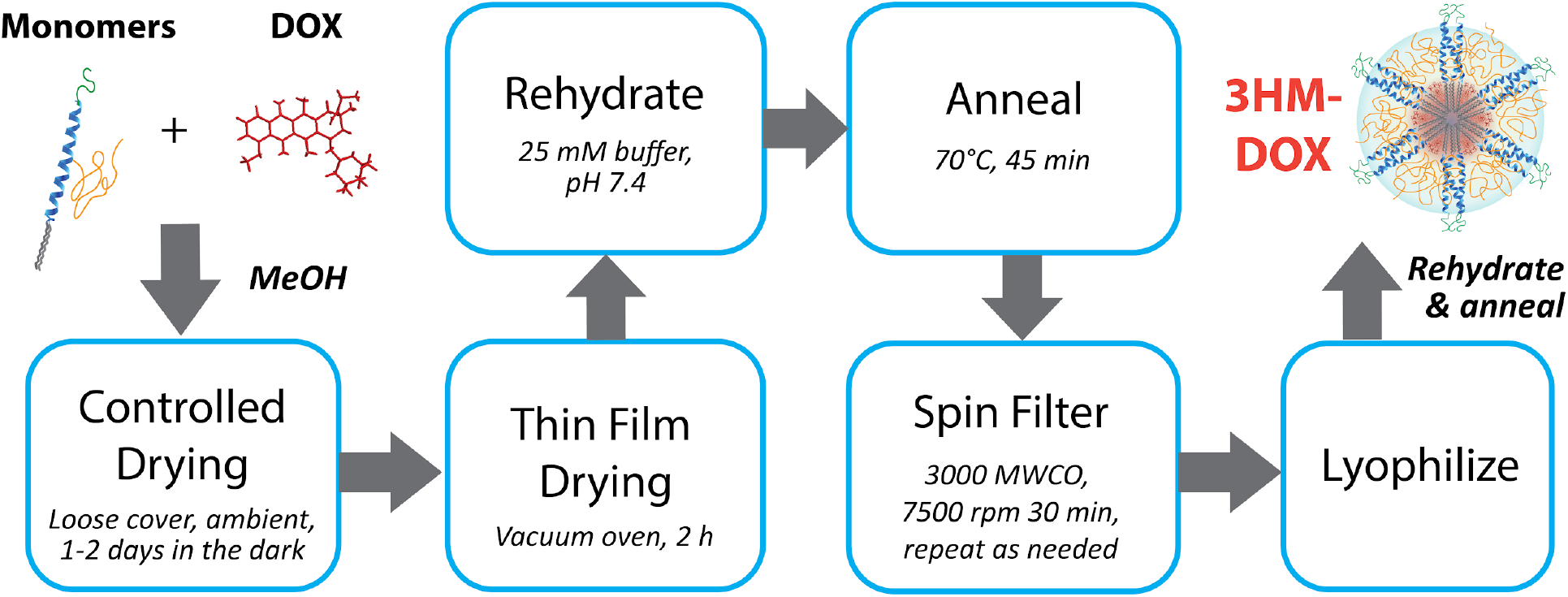
3HM-DOX cargo co-assembly process via thin film rehydration and self-assembly. In a similar fashion, other small molecule drugs (apomorphine, rapamycin, tamoxifen, and paclitaxel) were co-assembled in 3HM.

### 2.3: Drug Property Analysis

LogP values used as a quantitative measure of hydrophobicity were obtained from DrugBank. Hansen solubility parameter accounting for non-polar and Van der Waals interactions, δ_d_ was calculated by group contribution method. ^41^ Drug molecules are divided into smaller functional groups and the contributions from different groups were accounted together to evaluate δ_d_. Molecular dimensions of drug molecules were obtained from Chem3D Pro19.0. Energy minimized 3D structure of molecule were obtained by MM2 simulation function, designed to reproduce equilibrium molecular covalent geometry of molecules. ^42^ Subsequently, triplicate obelisk geometric measurements (detailed further in the main text) were made along a fixed axis for each molecule, and the mean dimensions are reported.

### 2.4: Small angle X-ray Scattering (SAXS)

Small-angle x-ray scattering (SAXS) experiments were carried out at the Advanced Photon Source (APS) at the Argonne National Lab, Argonne, Illinois at the 8-ID-I beamline. The instrument was operated using an X-ray energy of 10.9 keV, a sample-detector distance of 1.3 m, and a 1 M Pilatus detector. This provides an effective range of momentum transfer, Q, of 0.02 to 0.3 Å^−1^, where Q = 4π sin θ/λ, θ = scattering angle, and λ = 1.14 Å wavelength of incident X-rays. Samples were contained in standard boron–quartz capillaries situated in a homemade sample holder. Using this setup, background subtraction could be performed quantitatively. 3HM and 3HM-DOX samples were dissolved in phosphate buffer (25 mM, pH 7.4) at a concentration of ~5 mg/ml, annealed at 70 °C for 1 h and allowed to equilibrate at room temperature overnight before filtering with a 0.2 μm nylon filter (Pall). SAXS measurements were performed with 5 seconds acquisition times with no observable beam damage.

### 2.5: Small-Angle Neutron Scattering (SANS)

SANS experiments were conducted at beamline NG3 at the National Institute of Standards and Technology (NIST, Gaithersburg, MD) ^43^ using cold neutrons (6 Å) with two detector distances (5 and 13 m) to cover an effective range of momentum transfer, Q = 4π sin θ/λ (θ is the scattering angle, and λ is the wavelength of incident neutrons), from 0.013 to 0.4 Å^−1^. Both 3HM and 3HM-DOX samples with C16 hydrogenated (−(CH_2_)_15_CH_3_) and deuterated (−(CD_2_)_15_CD_3_) alkyl tails were prepared in a 25 mM KH_2_PO_4_ deuterated (D_2_O) buffer at pH 7.4. The amphiphiles do not include PEG750, to be directly comparable to prior SANS studies. ^44^ Samples were annealed at 70 °C for 1 h, and allowed to equilibrate at room temperature overnight before conducting SANS measurements. SANS data were normalized by the concentration of micelles (~7 - 20 mg/ mL). The SANS scattering intensity profiles were reduced to absolute scale with the NIST data reduction macros in IGOR Pro available from NIST. ^45^

### 2.6: Differential Scanning Calorimetry (DSC) and Kissinger Analysis

DSC Thermograms were obtained for buffered 3HM and 3HM-DOX solutions (~1 mg/mL in phosphate buffer, 25 mM, pH 7.4) using a VP-Microcal calorimeter (GE). ~550 μL samples were loaded and equilibrated at 5 °C for 15 min. Temperature was increased to 70 °C at 1°C/min scan rate. Thermograms were baseline corrected, normalized for exact concentration, and fit using Origin software provided with the VP-Microcal instrument to provide T_m_ peak maxima. For the Kissinger Method analysis, scan rates were varied (0.1 to 1.5 °C/min) in order to allow kinetic analysis. ^46^ Kinetic parameters can then be found by fitting values in an Arrhenius-type relationship that takes the form of Y = mX +b:

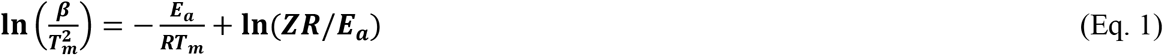

Where β is the scan rate, T_m_ is the measured melting temperature (point of iso-conversion), E_a_ is the kinetic activation energy, R is the universal gas constant, and Z is the energetic exponential prefactor. Kinetic parameters are related to the thermal transition of the alkyl chain core.

### 2.7: Fluorescence Recovery and Activation Energy of DOX Desorption

Fluorescence spectroscopy was performed on 3HM-DOX solutions using an LS-55 fluorescence spectrometer (PerkinElmer). Stock solutions of 3HM-DOX were prepared, aliquoted, and lyophilized for each experiment so samples had precisely the same concentrations and thermal histories. The test solutions were rehydrated to 0.2 mg/mL in phosphate buffer (25 mM, pH 7.4) and annealed at 70 °C for 45 min, then allowed to equilibrate at room temperature for 1 h before fluorescence recovery experiments. Samples were loaded into a 1 mm path length quartz cell (Starna Cells). Emission spectrum were recorded from 490 nm to 640 nm using an excitation wavelength of 450 nm at a scan rate of 200 nm/min. Temporal evolution of fluorescence intensity was recorded every 15 min at 522 nm. Temperature was controlled with a Peltier temperature controller (PTP-1, PerkinElmer). Fluorescence recovery experiments were conducted over 24 h for a range of temperatures, from 35 - 70 °C. Fluorescence intensity was normalized to initial fluorescence at time zero and fully released. 100% DOX release fluorescence intensity was obtained by dissolving the 3HM-DOX in methanol.

The resultant fluorescence recovery curves showed a sigmoidal behavior for scans taken at ≥ 45 °C. Since 3HM’s crystallized alkyl tail cores have a melting temperature ~ 40 °C, ^5^ the sigmoidal shape was assumed to include an initial melting stage followed by a slow DOX diffusional release stage. To derive E_a_ of DOX release, only data points past the inflection point of each series (for 55 - 70°C curves) were considered for calculations using the Arrhenius Equation. Rate constants from the negative exponential curves of % DOX remaining over time were plotted as ln(k) vs. 1/RT (kJ^−1^ mol) and fit to yield the E_a_ of DOX desorption.

### 2.8: Dialysis and Drug Release Kinetics Model

0.5 mL of 2 mg/mL 3HM-DOX (9.5% DOX content; 0.19 mg/mL DOX) was loaded into dialysis tubes (ThermoFisher, CAT# 88400, 3500 Da MWCO) and dialyzed against 15 mL of 25 mM phosphate buffer pH 7.4 (matched osmolality to 3HM-DOX formulations). Triplicate samples were prepared and incubated at 37 °C during the release experiment. Prior to the experiment, and at desired time-points, solution UV-Vis absorbance at 280 nm and 480 nm was measured by Nanodrop (4 uL droplet, 0.1 cm path length). The droplet was recovered and replaced in the sealed sample reservoir after each measurement. DOX concentration was determined using A480 (e = 14.509 cm^−1^ mL mg^−1^). The reservoir was replaced with fresh buffer daily. 3HM content was assumed constant with negligible diffusion through the dialysis membrane since monomer MW is ~7300 Da. Evaporative volume corrections were used for DOX concentration determination, where [DOX]_corr._ = [DOX]_meas._ * V_meas._ / 0.5 mL.

Detailed derivations of the drug release kinetics are provided in the **Supplementary Methods**. This analysis is based on a similar study done for block co-polymer micelles, ^47^ but without terms describing cleavage of covalently-bound drugs. For a dialysis system in non-infinite sink conditions, DOX release from 3HM is driven by free DOX diffusion through the dialysis membrane. This is represented by the rate equation: 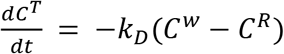, where *k*_*D*_ is the reaction rate constant, *C*^*w*^ is DOX concentration in the solvent (water) inside the dialysis chamber, and *C*^*R*^ is DOX concentration in the reservoir. A partition coefficient is assumed for DOX preference of the micelle over the bulk solvent, *K*_*p*_ = *C*^*m*^/*C*^*w*^, as illustrated in **Scheme 3**. A version of derivations assuming two partition coefficients (accounting for DOX preference in core/shell and shell/solvent) is included in the **Supplementary Methods**.

**Scheme 3:**
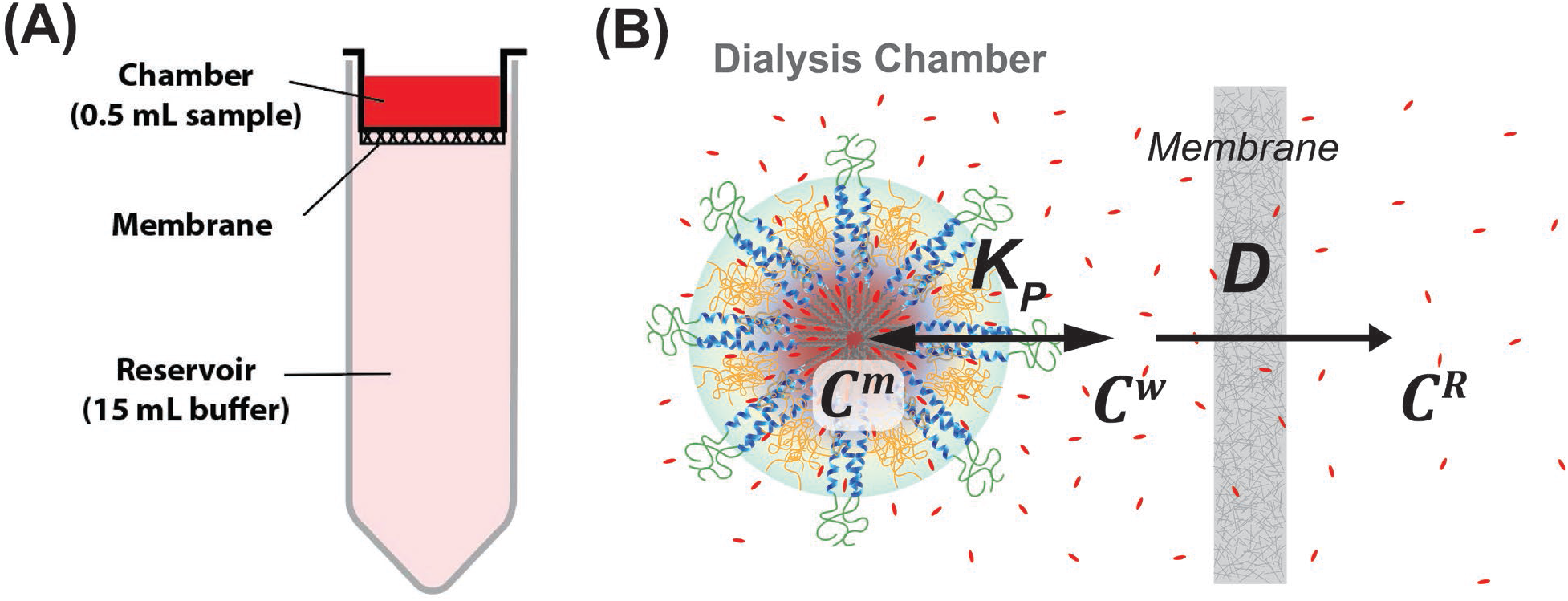
Schematic for (A) 3HM-DOX dialysis setup, and (B) DOX diffusion through the dialysis membrane to the reservoir, whereby DOX partitions between the micelle and surrounding solvent in the dialysis chamber.

Accounting for mass balance between DOX amounts in the micelle core, shell, and solvent, as well as the partition coefficients, yields the following pair of ordinary differential equations for flux of DOX moving out of the dialysis chamber to the reservoir:

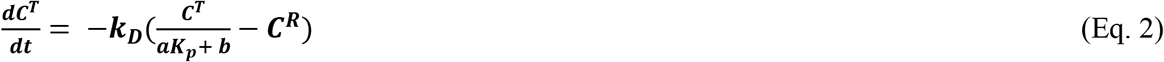

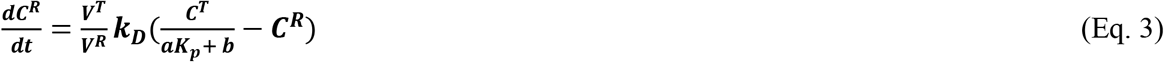

Volume fractions are captured in the following terms: *α* = *V*^*m*^/*V*^*T*^, and *b* = *V*^*w*^/*V*^*T*^, where *V*^*m*^ is the micelle volume, *V*^*w*^ is the surrounding solvent (water) volume inside the dialysis chamber, and *V*^*T*^ is the total volume of the dialysis chamber. *V*^*R*^ is the volume of the reservoir. *V*^*T*^ for 3HM-DOX solution volume is fixed at 0.5 mL. While the reservoir was changed daily (15 mL fresh PBS each time), to simplify the model fit, reservoir volumes were set as a constant (i.e. *V*^*R*^ = 15 mL for fitting single-day dataset (T = 0-13 h), and 30 mL for fitting two-day dataset (T = 0-36 h)).

Dialysis data (*C*^*T*^= DOX concentration in the dialysis chamber over time) were fitted to Eq. (2)–(3) using MATLAB to solve for *K*_*p*_ and *k*_*D*_. Boundary conditions at *t* = 0 for these equations are: 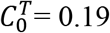 mg/mL, and 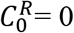mg/mL.

### 2.9: Spin Filtration to Determine 3HM-DOX Core-Shell Partition Coefficients

Three 3HM-DOX solutions, starting at 11 w/w% DOX, were prepared for spin filtration. 2 mg of 3HM-DOX (2 mL of 1 mg/mL solutions in 25 mM phosphate buffer pH 7.4) were loaded into 4 mL spin filter tubes (Amicon Ultra, 3000 MWCO). Samples were centrifuged at 7500 rpm for 30 mins. The filtrate was discarded, and the ~0.1 mL retentate was mixed with Milli-Q water to a total of 4 mL. After each spin, 5 μL was sampled and diluted 20x with MeOH for triplicate UV-Vis measurement at A280 and A480. For Tube 1, spin filtration was repeated 3x, then the sample was lyophilized. Tube 2 was spin filtered 6x before lyophilization. Tube 3 was spin filtered 8x, then stored at 5°C in the dark overnight. UV-Vis measurement was taken before and after Tube 3 storage to verify A480 and A280 ratios had remained unchanged. Tube 3 sample was spin filtered 8x more on Day 2, before lyophilizing.

The ratios of A480/280 values were tracked as a function of the number of spins to approximate DOX removal relative to 3HM peptide retained. It would have been difficult to determine 3HM DOX % loading by UV-Vis alone during the experiment, since DOX UV absorbance at A280 overlaps with the peptide signature. As such, samples were removed at spin #3 (Tube 1), #6 (Tube 2), and #16 (Tube 3) for lyophilization and weighing of the 3HM-DOX powder. After re-dissolving the powder in buffer and taking A480 measurements (DOX **ε** = 22.5 cm^−1^mg/mL^−1^, see **Supplementary Figure S1** for DOX calibration curve), DOX % loading was obtained from the known 3HM-DOX masses. As the ratio of A480/280 reached a plateau after multiple spins, loosely-bound DOX has been presumably removed from the 3HM shell. Using the difference in DOX % loading at the beginning and after the plateau, the DOX partition coefficient between 3HM core to shell were calculated.

## 3. Results and Discussion

### 3.1: Importance of Geometry and Hydrophobicity for Cargo Loading

For sub-20 nm nanocarriers, consideration should be given to small molecule cargo impact on the overall assembly. Loading content in micelles generally correlates with cargo hydrophobicity or LogP values, with higher loading for more hydrophobic drugs. ^48–50^ As the nanocarrier size approaches that of its cargo, however, the interplay between cargo molecular structure and amphiphile packing parameter becomes more significant.

To delineate the effect of hydrophobicity and molecular geometry, therapeutic small molecules with established clinical relevance were tested: doxorubicin (DOX), ^51^ apomorphine (APO), ^52^ rapamycin (RAP), ^53^ tamoxifen (TAM), ^54,55^ Dexamethasone (DEX), ^56^ and paclitaxel (PAX). ^57^ Hansen solubility parameters^41^ based on non-polar interactions, δ_d_, were calculated for drugs (~19-21 (J/cm^3^)^0.5^ as shown in **Table 1**) versus micelle core component C18 stearic acid (17.9 (J/cm^3^)^0.5^). Interestingly hydrophobic interactions alone did not account for the compatibility of the tested drugs with the 3HM core. **Figure 1A** shows no obvious correlation between the LogP values of these drugs (ranging from ~2 - 6) and their loading efficiency.

**Table 1:** Physicochemical properties of the drugs that were investigated for their formulation in 3HM. Loadings obtained for each drug represent average from 5 different measurements.

| Drug | Log P | Molecular weight (Da) | Solubility parameter, $\delta_d$ (J/cm<sup>3</sup>)<sup>0.5</sup> | Loading (% w/w) |
| --- | --- | --- | --- | --- |
| Tamoxifen (TAM) | 5.93 | 371 | 19.11 | 1.5±0.3 |
| Rapamycin (RAP) | 4.85 | 914 | 20.29 | 3.3±0.2 |
| Paclitaxel (PAX) | 3.21 | 853 | 21.15 | 2.5±0.3 |
| Apomorphine (APO) | 2.51 | 267 | 19.29 | 7.6±0.5 |
| Dexamethasone (DEX) | 1.93 | 392 | 20.37 | 1.8±0.2 |
| Doxorubicin (DOX) | 1.91 | 543 | 20.06 | 7.9±0.4 |
| Stearic acid | 8.23 | 284 | 17.90 | N/A |

**Figure 1:**
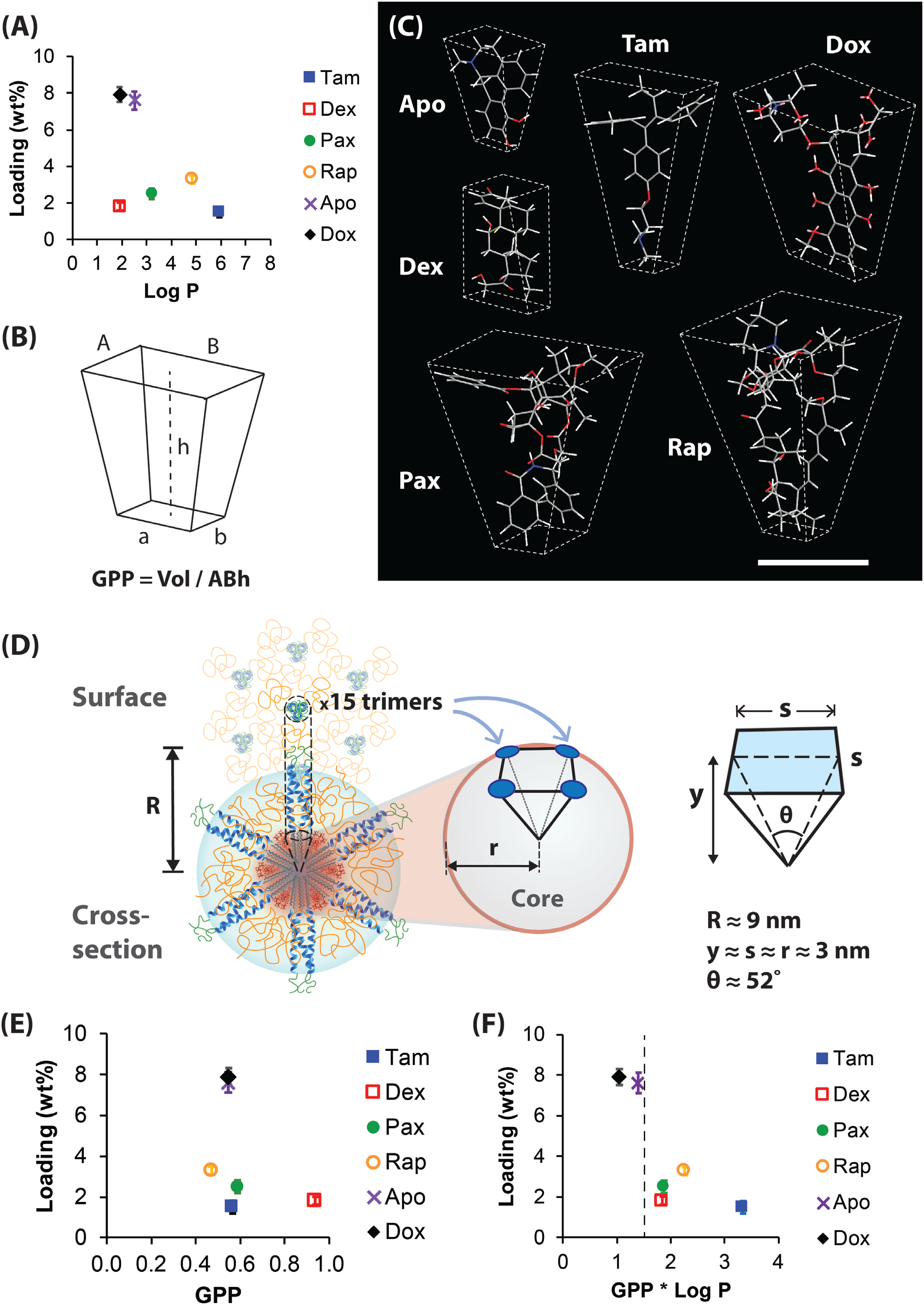
(A) Drug loading shows no correlation to LogP hydrophobicity index. (B) Obelisk shape dimensions used for calculating the geometric packing parameter, GPP. (C) Three dimensional structures of drug molecules generated in Chem3D, fitted to obelisk shapes (scale bar = 10 angstroms). (D) Schematic of 3HM core divided into ~13 pyramidal shapes with an amphiphilic trimer at each vertex. (E) Drug loading shows no correlation to GPP alone, but (F) correlates with the product of GPP and LogP values, with high drug loading at GPP*LogP < 1.5.

Presumably, geometric packing between small molecules and nanocarrier amphiphiles plays a critical role in determining cargo loading compatibility, akin to amphiphile critical packing parameter (CPP) and self-assembly architecture. ^18^ The 3HM core radius (3 nm) ^44^ is comparable to the ~1 nm molecular dimensions of drug molecules studied here, assessed by Chem3D (Cambridge Soft) using the built-in MM2 simulation function. ^42^ We hypothesize that wedge-like or planar drugs can intercalate among the C18 tails in 3HM core (**Scheme 1A**) without compromising alkyl tail crystalline packing, unlike bulky drugs with isotropic geometry.^35,37,39^

Each drug molecule was approximated as a trapezoidal obelisk with asymmetrical top and bottom planes (see **Figure 1B** and **1C**). To quantify the degree of structural asymmetry, we propose a “geometric packing parameter” (GPP), analogous to the calculating the CPP,^18^ defined by Equation 4. The GPP obelisk volume is given by 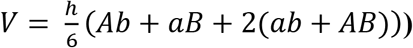. Here, *AB* is the dimensions of the larger top face, *ab* is the smaller bottom face, and *h* is the height of the obelisk. A GPP value closer to 1 indicates the drug is approximately rectangular (cuboid) in shape, while a low GPP value indicates a more pyramidal structure.

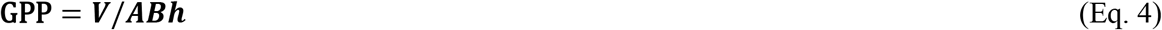

The obelisk dimensions and GPP value will determine whether a molecule can fit into the wedge-like spaces between 3HM core alkyl tail bundles (shown in **Scheme 1C**). Since each 3HM particle is comprised of ~15 trimeric bundles (45 monomeric subunits) ^37^, the alkyl tails are predicted to create inverted pyramidal spaces in the core (**Figure 1D**), roughly equivalent to a 13-face polyhedron. For a sphere with 3 nm radius (surface area 113 nm^2^, volume 113 nm^3^), each pyramidal section has ~3 nm-sided base with 8.7 nm^2^ surface area, and 8.7 nm^3^ volume. Thus, each inverted pyramid has a small GPP value of 0.33. As shown in **Table 2**, all drug GPP values exceed 0.33, but since the largest dimension among them is ~2 nm, all of them would fit within these pyramidal sub-spaces.

**Table 2:** Results of geometric analysis of drug molecules modeled in Chem3D. Dimensions reported are the average of three measurements per molecule.

| Drug | Loading<br>(% w/w) | A<br>(nm) | B<br>(nm) | a<br>(nm) | b<br>(nm) | h<br>(nm) | V<br>(nm<sup>3</sup>) | GPP | Log<br>P | GPP×<br>LogP |
| --- | --- | --- | --- | --- | --- | --- | --- | --- | --- | --- |
| Tam | 1.5±0.3 | 1.1 | 0.4 | 0.3 | 0.3 | 1.5 | 0.409 | 0.56 | 5.93 | 3.34 |
| Dex | 1.8±0.2 | 0.6 | 0.5 | 0.7 | 0.4 | 1.1 | 0.333 | 0.96 | 1.93 | 1.85 |
| Pax | 2.5±0.3 | 1.3 | 0.9 | 0.5 | 0.6 | 1.5 | 1.079 | 0.58 | 3.21 | 1.87 |
| Rap | 3.3±0.2 | 1.9 | 1.1 | 0.5 | 0.4 | 1.8 | 1.637 | 0.47 | 4.85 | 2.26 |
| Apo | 7.6±0.5 | 0.7 | 0.4 | 0.4 | 0.2 | 0.9 | 0.156 | 0.56 | 2.51 | 1.40 |
| Dox | 7.9±0.4 | 1.0 | 0.9 | 0.5 | 0.4 | 1.5 | 0.690 | 0.55 | 1.91 | 1.05 |

Ultimately, the data confirmed that for 18 nm 3HM, the drug loading efficiency is not determined by the LogP nor GPP alone **(Figure 1E)**, but by the product of LogP and GPP (**Figure 1F**). When GPP*LogP is greater than 1.5, drug loading drops sharply from ~8% to < 4% w/w. As such, higher loading requires a combination of both low GPP (more pyramidal) and low LogP values (less hydrophobic; LogP > 0). The implications from low LogP values is surprising, since “like dissolves like” compatibility is expected between small molecules and the hydrophobic C18 alkyl tails. The results suggest that maximal drug packing occurs at a different microenvironment, such as the interface between the hydrophobic core and the more hydrophilic peptide-polymer shell. This is explored with SAXS and SANS measurements of the internal structure of 3HM-DOX, shown in the next section.

Also notably, DOX and APO appeared to have the most planarity in comparison to other drugs tested, when viewed from the angle of their thinnest side (**Supplementary Figure S2**). A flat geometry would be compatible with molecular packing among crystallized alkyl chains, as discussed in the next section. Additionally, while the drug simulation method chosen for these experiments (MM2 function in Chem3D) is rather “rudimentary” compared to cutting-edge Monte Carlo or molecular dynamics methods (e.g. Large-scale Atomic/Molecular Massively Parallel Simulator (LAMMPS)), these MM2 simulations are much more accessible to many researchers, and still provide insightful analysis in the context of other knowledge. For 3HM alone, particle structure generated by molecular dynamics has been previously reported.^44^

Overall, the results presented in this section underscore the significance of studying atomic-scale physicochemical features of cargo molecules, which is not commonly investigated in detail for micellar loading capacity. GPP and hydrophobicity analysis can be applied to other small nanocarrier systems (e.g. block co-polymer micelles), whereby the specific internal structures of these systems must be considered.

### 3.2 3HM-DOX Internal Structure

#### 3.2.1 DOX Distribution via SAXS and SANS

To molecularly characterize the distribution of DOX within 3HM, and understand any structural changes induced by DOX co-assembly, small angle X-ray and neutron scattering (SAXS, SANS) were employed. ^31,32^ Understanding which components of 3HM directly interface with DOX is useful to later modulate DOX release behavior overall nanocarrier stability, and eventual *in vivo* performance. Investigating the structural details of small molecule drug and small nanocarrier interactions also provides experimental evidence supporting the geometric compatibility analyses presented earlier.

While they require significant resources and analysis, SAXS and SANS are advantageous in probing the internal structure of soft nanomaterials over typical methods such as dynamic light scattering (DLS) and transmission electron microscopy (TEM). ^31^ Contrast in these scattering techniques originates from differing scattering length densities (SLDs) of the individual components within a system, shown for 3HM(-DOX) in **Supplementary Table S1** and **S2**, for X-rays and neutron studies respectively. ^58–61^ This is schematically illustrated in **Figure 2A**. Particular to SANS, contrast variation by selective deuteration (based on SLD differences in hydrogen and deuterium) enables simultaneous fitting of multiple contrast measurements. This has been demonstrated with regards to PEG2000 location and water content found in 3HM. ^44^ Here, selective deuteration of the alkyl core allows the internal dimensions of 3HM to be highlighted, allowing the interpretation of structural changes induced by DOX incorporation, using prior SAXS and SANS studies of unloaded 3HM as a foundation. ^4,37,44^

**Figure 2:**
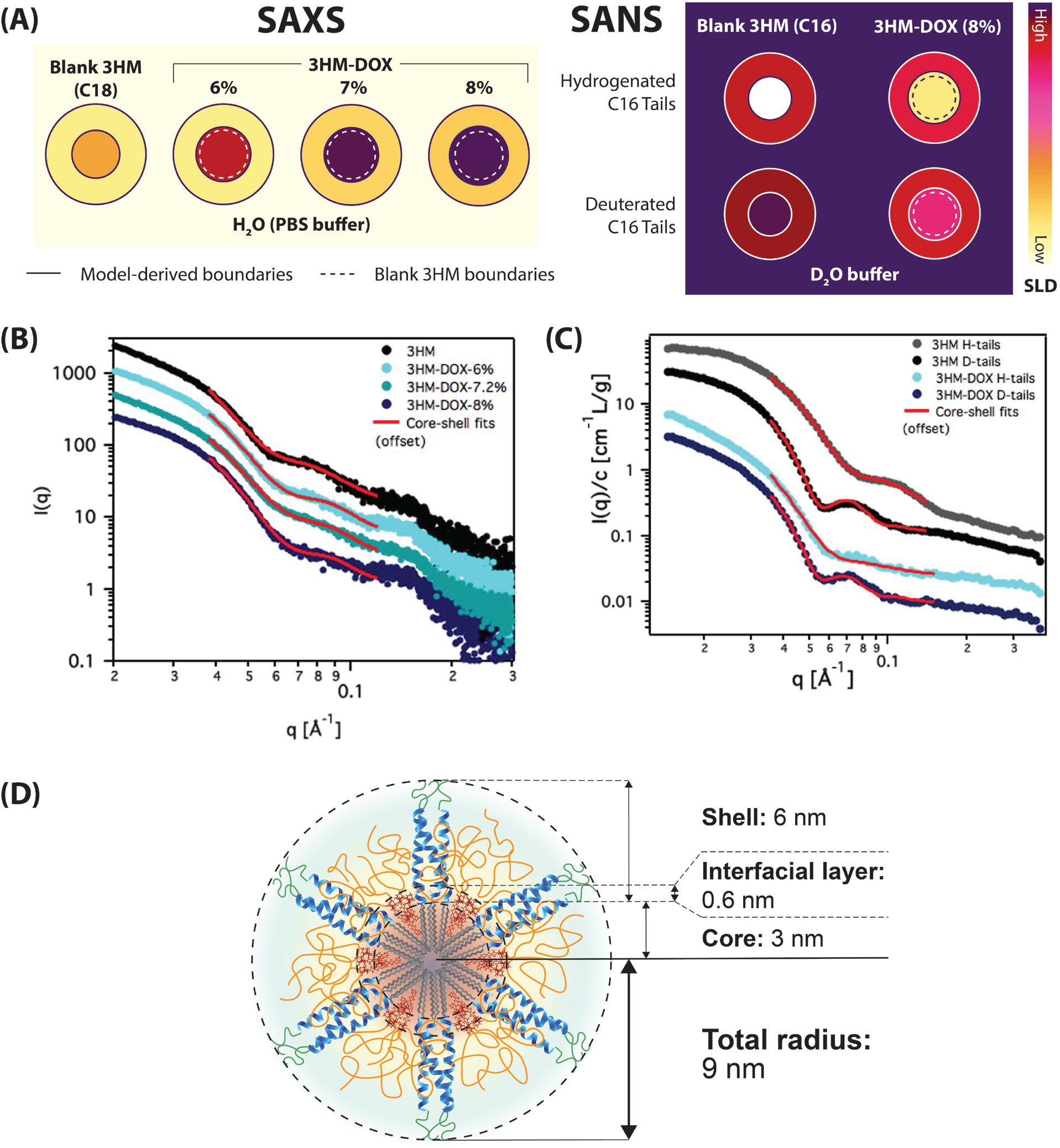
SAXS and SANS analysis of 3HM-DOX internal structure. (A) Schematic of core-shell SLD contrasts and boundaries derived from the analysis: in SAXS, the analysis is done in water, while in SANS, the analysis is done in deuterium solvent. (B) SAXS intensity profiles and core-shell (sphere) fits of 3HM (C18) with increasing DOX loading. (C) SANS intensity profiles and core-shell (sphere) fits of 3HM (C16, without P750) and corresponding 3HM-DOX 8% w/w. C16 tails were either hydrogenated (H) or deuterated (D). Profiles are offset for clarity. (D) Idealized schematic of 3HM-DOX with core-shell thicknesses obtained from SAXS fitting of unloaded 3HM(C18), with an interfacial layer occupied by DOX at 8% w/w loading.

SAXS studies investigated a variant of 3HM (with C18 tails and PEG750 surface) co-assembled with three different loadings of DOX (**Figure 2 A** & **B**). This 3HM variant has clinical potential based on its physical stability, long *in vivo* circulation, and sufficient core volume to accommodate DOX. ^4^ X-ray scattering intensity profiles were fit with core-shell models, allowing polydispersity to account for mobile PEG side-chains being able to extend past the peptide ^39^ (see **Scheme 1** for 3HM monomer components).

At higher DOX loading, 3HM core SLD increased accordingly while the shell SLD stayed relatively constant (**Supplementary Table S3**). This indicates DOX localizing within the 3HM alkyl tail core, while the peptide shell layer and outer surface remain relatively free of DOX. The core radius increased 0.3 nm with DOX loadings of both 6% and 7.2% w/w, with a further increase of 0.3 nm for 8% DOX, at the apparent cost of shell thickness. The overall micelle size remained constant regardless of increasing DOX content, in agreement with 3HM-DOX DLS and TEM characterization reported previously (**Table 3**). ^5,38^

**Table 3:** Fitted dimensions obtained by SAXS for 3HM and 3HM-DOX components

| **Sample** | **3HM**<br>**(C18)** | **3HM-DOX**<br>**6 %** | **3HM-DOX**<br>**7.2 %** | **3HM-DOX**<br>**8 %** |
| --- | --- | --- | --- | --- |
| Core radius (nm) | 3.0 | 3.3 | 3.3 | 3.6 |
| Shell thickness (nm) | 6.0 | 5.7 | 5.7 | 5.4 |
| Total radius (nm) | 9.0 | 9.0 | 9.0 | 9.0 |

Presumably, the observed internal dimension changes stemmed from DOX locating at the core-shell interface **(Figure 2D).** The assembly of 3HM’s 1CW alpha-helical peptide headgroup stays robust upon DOX addition, ^38^ such that the peptide shell dimensions likely remain unchanged (i.e. peptide does not unfold). Conceivably, there is higher free volume toward the outer core due to the curvature of 3HM micellar assembly, as shown in **Scheme 1C**. Corresponding with the geometric analysis presented earlier, DOX would first occupy all the available free volume in the core, then invade radially outward towards the core-shell interface.

This allows the maintenance of 9 nm overall radius, regardless of increasing DOX content. If DOX had instead accumulated within the core’s center, alkyl rearrangement and a corresponding increase in total micelle radius would have been expected. However, this was not observed.

The interfacial DOX layer is likely accommodated as part of the core during fitting due to SLD similarity (**Figure 2A)**. SAXS results are fitted to a two-layer (i.e. core-shell) model, and DOX has relatively high SLD compared to the shell components (**Supplementary Table S1)**. With the maintenance of overall size, these observations support the location of DOX to the core-shell interfacial space, without deleterious effects on 3HM internal structure, uniform size, and surface properties.

To enhance contrast of the internal components of the micelle core, SANS studies were conducted with selective deuteration of the alkyl tails. A 3HM variant with C16 alkyl tails and without PEG750 was investigated for direct comparison against prior SANS studies with unloaded 3HM (C16). ^44^ 3HM and 3HM-DOX samples prepared with hydrogenated (H) or deuterated (D) alkyl tails were analyzed in D_2_O buffer solutions (**Figure 2A** & **C** and **Supplementary Table S4**). Simultaneous fits provided core-shell 3HM dimensions shown in **Table 4**. In comparison to the SAXS 3HM(C18) dimensions shown above, the core radius was smaller (2.7 vs. 3 nm) corresponding with decreased tail length, and the shell was smaller (5 vs. 6 nm) due to the omission of outermost-layer PEG750. ^44^ DOX (~8 % w/w) introduction increased the thickness of the core by ~0.2 nm at the cost of shell thickness, consistent over both core contrasts. Unlike with SAXS, both the 3HM(C16) core and shell SLD changed upon DOX incorporation. DOX SLD for SANS is higher than that of C16, peptide and PEG components, but lower than that of D_2_O or deuterated C16 (**Supplementary Table S4**). Due to high D_2_O content in the 3HM shell, its SLD is similar to the deuterated core and contrasts against the hydrogenated core. The corresponding SLD changes for 3HM(C16)-DOX is thus consistent with DOX incorporating into both the core and shell layers, while keeping the overall particle radius relatively constant (**Figure 2A**).

**Table 4:** Fitted values for SANS for C16 3HM and 3HM-DOX components.

| Sample | 3HM (C16) | 3HM-DOX (C16) |
| --- | --- | --- |
| Core radius (nm) | 2.7 | 2.9 |
| Shell thickness (nm) | 5.2 | 5.0 |
| Total radius (nm) | 7.8 | 7.9 |

The SANS results support the development of a DOX-rich interfacial layer between the core and the shell, similar to SAXS of 3HM(C18)-DOX. Since C16 tails have a correspondingly smaller core volume than C18 tail micelles, they are unable to fully accommodate DOX without slightly expanding (by 0.1 nm) the overall micelle size (**Table 4**). The 3HM(C16)-DOX core SLD of both hydrogenated and deuterated contrasts corresponded to ~29 v/v% DOX in the core, or ~9 % w/w overall, in close agreement with the 8 % w/w determined by UV-vis. Importantly, these scattering studies confirm that 3HM co-assemblies with DOX maintain their uniform overall size and shape. Being reliant on fitting to constrained models, there is some space for interpretation of exact dimensions in isolated SAXS and SANS results. It is important to reiterate that these conclusions align with previous scattering, TEM, and DLS data, ^4,5,37,38,44^ and correlate well with release and thermal behavior of 3HM-DOX co-assemblies developed subsequently in this contribution. Combined with following experiments, showing no burst release upon dissolution, they also indicate that DOX is protected from the surface of the micelle. Further, DOX locating towards the core-shell interface of 3HM affirmed the implications of optimal cargo hydrophobicity and geometry shown earlier.

#### 3.2.2 Alkyl Tail Rearrangement Upon DOX Co-assembly

Having detailed the spatial distribution of DOX within 3HM core and interface, the effect of DOX co-assembly on alkyl tail arrangement was examined. DSC measurements showed that the alkyl tail melting temperature (T_m_) (indicative of the degree of lipid crystallinity, ^33^) of 3HM(C18)-DOX (8 % w/w) is ~8°C higher than the 30°C T_m_ for unloaded 3HM. ^38^ The enthalpy (∆H) change is 33.5 kJ/mol and 25.3 kJ/mol for unloaded and loaded micelles, respectively.

3HM(C18)-DOX alkyl tail T_m_ increased with DOX loading, as shown in **Figure 3** and **Table 5**. This implies increased alkyl chain crystallinity with DOX incorporation, albeit still remaining below 100% crystallinity C18 (octadecane T_m_ = 57°C). ^62^ The observed T_m_ shift is proposed to arise from the specific molecular arrangement adopted upon DOX co-assembly (**Figure 3**). Namely, the radiating splayed alkyl tails of unloaded 3HM are poorly ordered, with more free volume closer to the core-shell interface. DOX co-localization into these spaces would then enable alkyl tails rearrangement into more crystalline structures, owing to the favorable DOX GPP and planarity (discussed above in **Figure 1** and **Supplementary Figure S2**).

**Table 5:** Thermodynamic parameters gleaned from DSC curves of 3HM-DOX at various drug loading content.

| DOX Content<br>(% w/w) | $T_m$ ($^{\circ}\text{C}$) | $\Delta H$<br>(kJ/mol) | $\Delta S$<br>(kJ/mol$^{\circ}\text{C}$) |
| --- | --- | --- | --- |
| 0 | 29 | 46.4 | 1.63 |
| 6.8 | 30 | 39.1 | 1.38 |
| 7.3 | 32 | 35.5 | 1.11 |
| 9.0 | 37 | 34.2 | 0.96 |

**Figure 3:**
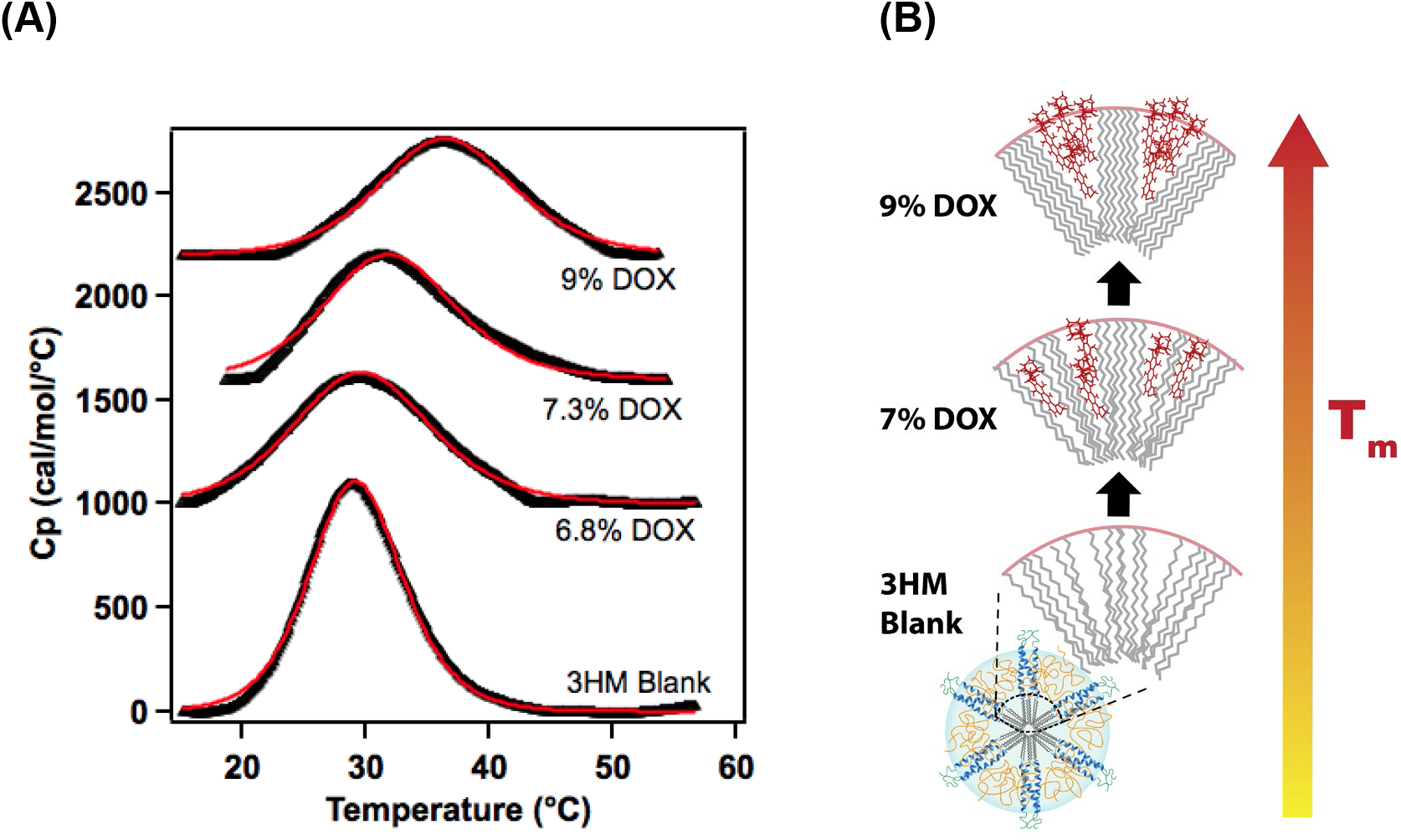
(A) DSC thermograms of 3HM-DOX at various loading content, showing exothermic peaks corresponding to the alkyl tail T_m_. (B) Idealized schematic of alkyl tail rearrangement to crystalline ordered structures at higher DOX loading, leading to higher T_m_.

Interestingly, the enthalpy (∆H) of alkyl tail melting decreased with DOX loading, in reverse of the T_m_ trend. From the DSC curves, the change in entropy (∆S) could be calculated via the relationship *ΔS* = *∫* (*C*_*p*_/*T*) *dT*. ^63^ Surprisingly, ∆S similarly decreased as a function of DOX loading (**Table 5**). A greater entropy change is expected from melting more well-ordered crystalline alkyl tails (i.e. breaking more intermolecular bonds). Interactions between DOX-C18 tails, and among crystallized DOX-DOX molecules potentially account for this behavior. At higher DOX loading, these interactions are more prominent and persistent throughout DSC scans. (thermograms were measured up to 70°C, while pure DOX crystals T_m_ = 230°C).^64^ Even though the alkyl tails had melted, the overall order within the core could remain high with these DOX-C18 and DOX-DOX interactions, which reduced the magnitude of ∆S compared to unloaded 3HM. Additional studies would be needed to confirm this theory. However, the overall results affirm that DOX co-localization increased 3HM core ordering.

### 3.3 Co-assembly Energy Landscape

With indications that cargo co-assembly can strongly affect nanocarrier architecture, corresponding impacts on subunit exchange and cargo desorption energy barriers were expected. To examine this, fluorescence (FL) experiments were conducted for Arrhenius analysis of DOX release, and DSC experiments were conducted for Kissinger analysis of 3HM core disassembly thermodynamics. Together, these methods revealed individual energetic contributions of DOX and the alkyl tails for the nanocarrier complex assembly.

#### 3.3.1: Activation Energy of DOX Release

FL release experiments are based on DOX molecules self-quenching while confined within the micelle core. As DOX is liberated into the bulk solvent (phosphate buffer), FL intensity increases. As shown in **Figure 4A**, there is negligible change in FL intensity at temperatures below the alkyl core melting transition (<46°C). This indicates significant interaction of DOX with the alkyl core, which confers high stability of 3HM-DOX in ambient conditions. Above the energetic barrier of the alkyl melting transition, 3HM subunits gained greater mobility, promoting sigmoidal DOX release at ≥ 45 °C. Prior to the inflection point, energy input is presumably required for transitional phase behavior of the alkyl tails, slowing release. Past the inflection point, DOX release profiles aligned closely with prior experiments conducted in bovine serum albumin (BSA) solution. ^5^ Note that for the current experiments, BSA was not included as a sink for amphiphile desorption^65,66^ in order to focus on 3HM amphiphile-DOX interactions.

**Figure 4:**
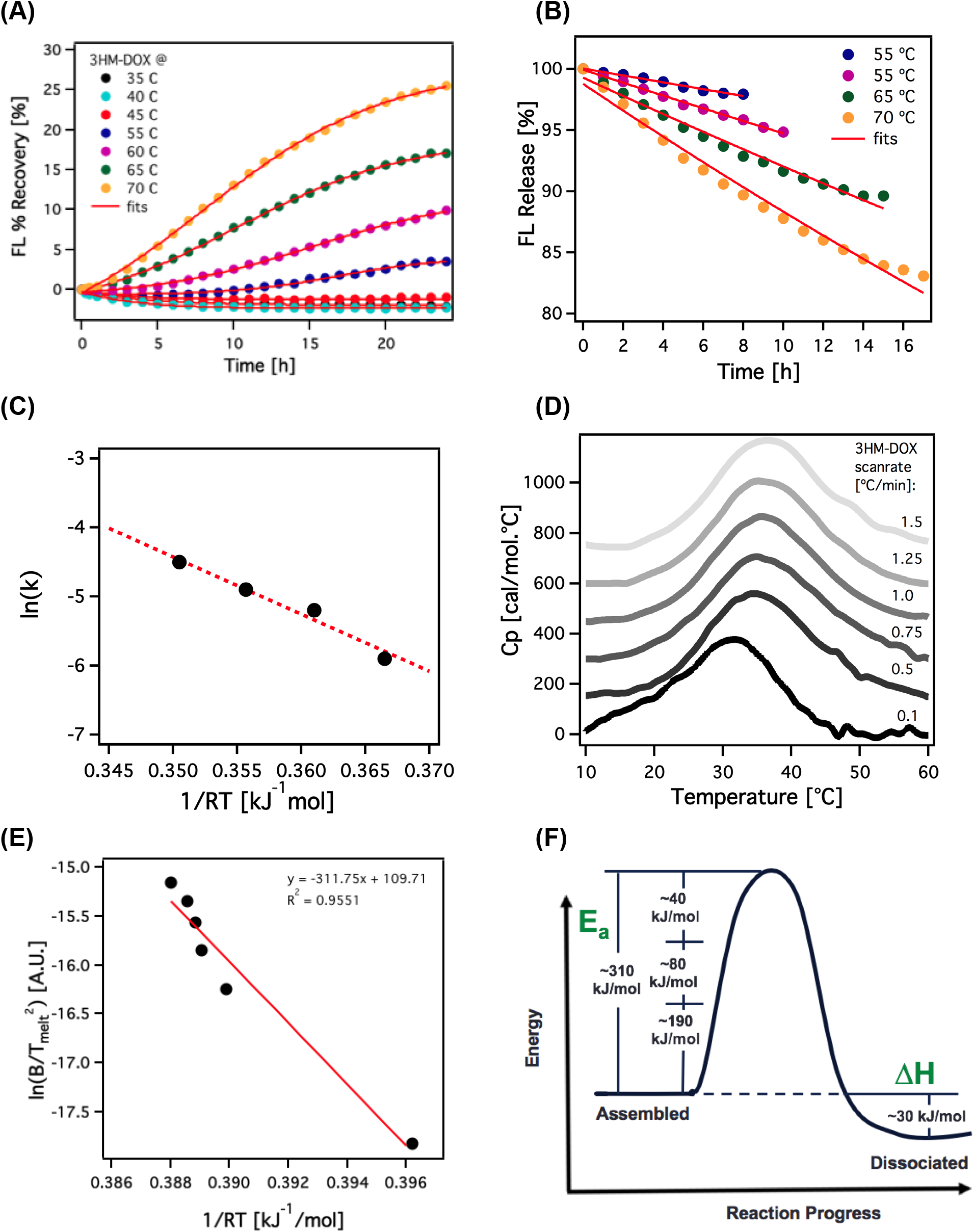
(A) FL release profiles of DOX from 3HM-DOX over temperature ranges. (B) Profiles truncated past the inflection point where fitting can be performed to Arrhenius-type release models. (C) Release constants can be plotted to yield an activation energy of 83 kJ/mol for the process of DOX release. (D) DSC thermograms of 3HM-DOX under different scan-rates to allow kinetic analysis of core alkyl tail-DOX interactions. (E) Kissinger kinetic analysis of DSC 3HM-DOX thermograms, yielding an activation energy of 312 kJ/mol. (F) Energy landscape of 3HM-DOX co-assembly including contributions of DOX and 3HM amphiphiles (monomer interactions ~190 kJ/mol, DOX-C18 ~80 kJ/mol, DOX-DOX ~40 kJ/mol) to the kinetic activation energy.

Using data from 55 - 70 °C scans past their inflection points, rate constants of DOX release were obtained, as shown in **Figure 4B** and **Supplementary Table S5**. Fitting these rate constants to an Arrhenius equation yields an activation energy of 83 kJ/mol for the process of DOX desorption, as shown in **Figure 4C**. This is comparable in magnitude to the 3HM subunit desorption activation energy, previously determined for 5,6-carboxyfluorescein (FAM)-labeled 3HM C16 to be 145 kJ/mol. ^38^ Considering the comparable magnitude of energy required for both processes, DOX release and subunit desorption likely occur past a similar critical temperature.

#### 3.3.2: Energetics of Overall 3HM-DOX Assembly

3HM-DOX core disassembly kinetics were probed by DSC to measure thermal interactions among DOX and the alkyl core. Subjecting 3HM-DOX to different thermal scan rates (**Figure 4D**), revealed the core energetic transitions of 3HM-DOX (**Supplementary Table S6**). Tracking the shift in T_m_ as a function of scan rate using Kissinger analysis allowed the activation energy of the transition to be calculated as 312 kJ/mol (Eq. 1; **Figure 4E**). ^46,67^ This quantity constitutes an apparent value, not to be interpreted as absolute truth. However, based on current molecular understanding, it is thought to embody energetic contributions from DOX-C18 tail, DOX-DOX, and C18-C18 desorption energies.

From previous kinetic analysis, it is known that 3HM with C16 tails display an E_a_ of 145 kJ/mol for monomer desorption. ^38^ ∆H of alkyl tail formation have also been reported as 25.1 and 33.5 kJ/mol for 3HM C16 and C18 tails respectively. ^38^ Assuming similar proportionality of E_a_ as with enthalpy for C16 vs. C18, the monomer desorption E_a_ for 3HM C18 can then be estimated as ~193 kJ/mol. As such, combining the kinetic E_a_ of DOX release (~83 kJ/mol; **Figure 4C**) and monomer desorption yields a total theoretical E_a_ of ~276 kJ/mol, which approaches the experimentally determined E_a_ of 312 kJ/mol. The remainder of the activation energy (~36 kJ/mol) may be attributed to DOX-DOX interactions, though further study is warranted for confirmation. Contributions of individual components to the system’s energy landscape is shown schematically in **Figure 4F**.

In sum, the 3HM-DOX energy landscape contains a very high kinetic barrier to molecular disassembly, despite having a ∆H on the scale of biological processes (e.g. ATP hydrolysis ~70 kJ/mol) ^68^. Geometric considerations discussed earlier are hypothesized to play a central role in this stabilization, with DOX intercalation into the alkyl tails near the interface alleviating the curvature induced by the bulky headgroup of 3HM. The magnitude of kinetic stabilization of DOX co-assembly was notable (~193 kJ/mol to 312 kJ/mol; a ~62% increase of E_a_,), exemplifying the consideration of the therapeutic cargo as an integral component of a co-assembly. These structural and thermodynamic insights are important for the ongoing development of frameworks to design and evaluate nanocarriers with precision, upon introduction to more complex environments.

### 3.4 3HM-DOX Partitioning and Release Kinetics

High energetic barrier of nanocarrier-cargo disassembly was expected to slow the cargo release kinetics. To quantify this, dialysis of 3HM-DOX formulations against phosphate buffer reservoir was performed (**Scheme 3**). As shown in **Figure 5A**, DOX content steadily decreased to ~55% of initial loading within ~40 hours. Fitting the release profile to the Korsmeyer-Peppas model^69^ (fractional DOX released, **f** = **kt**^**n**^) yielded **n** = **0.54** and **k**=**0.0685 h-0.54**(R^2^ = 0.9911) corresponding to **t_1/2_** = **40 h**. Negative exponential fitting yielded similar t_1/2_, further corroborating the 3HM-DOX slow release behavior **(Supplementary Figure S3)**. In the Korsmeyer-Peppas release model, Fickian diffusion corresponds to the value of n ≤ 0.45. ^70,71^ Hence, 3HM-DOX release fell outside of this regime, indicating other factors govern drug release beyond simple diffusion.

**Figure 5:**
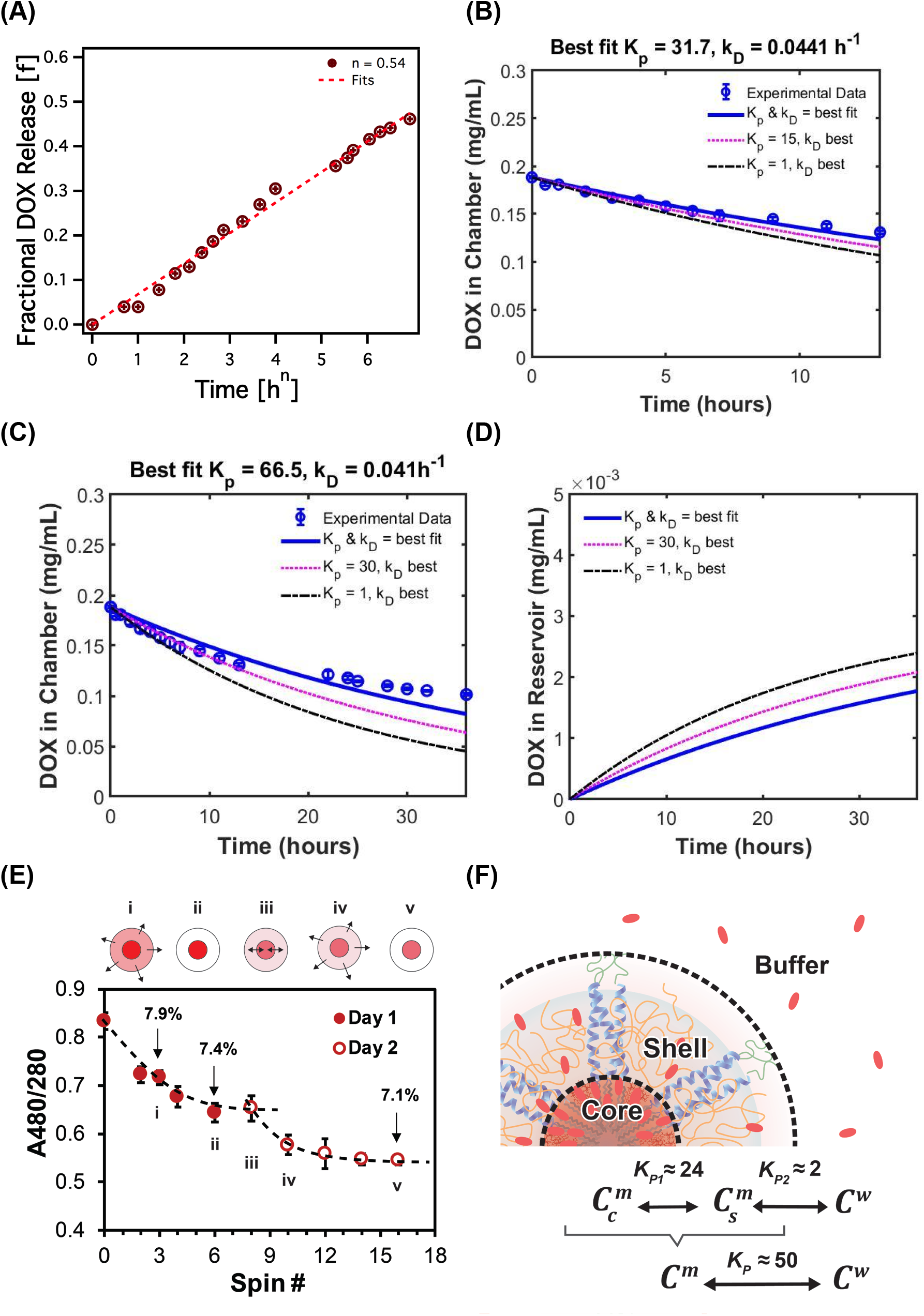
(A) Fractional DOX release fit to Korsmeyer-Peppas model: **f** = **kt**^**n**^ best fits (R^2^ = 0.9911) were found when n = 0.54, yielding k = 0.0685 h^−0.54^. (B) Results were fitted for T = 0-13 h (single-day) and (C) T = 0-36 h (two-days) datasets. Best-fit *k*_*D*_ and *k*_*p*_ values are shown for both, and trends with simulated *k*_*p*_ values are included. (D) Simulated concentration of DOX in reservoir, using various *k*_*p*_ values and *k*_*D*_ at 0.04 h^−1^. (E) Relative DOX (A480) to peptide (A280) UV-vis absorbance tracked after multiple spin filtrations over two days. Measured 3HM-DOX % w/w is shown for spin 3, 6, and 16. This process is schematically represented as (i) loosely-bound DOX being removed from 3HM shell during spins until (ii) only the core DOX remains, (iii) DOX re-partitioning between the core and shell during overnight storage, and (iv) additional DOX being removed from the shell on the next day until (v) a lesser amount remains in the core. (F) Schematic of DOX partition coefficients between 3HM core/shell (K_p1)_ and shell/bulk solvent (K_p2_), as well as overall DOX partition coefficient between 3HM/solvent, K_p_.

A system of rate equations involving drug partitioning (*K*_*p*_ for DOX in the micelle/bulk solvent) was derived, assuming diffusion across the dialysis membrane as the only driving force for cargo release (Eq. 2 and 3). With differing DOX co-localization in the core and shell of 3HM, *K_p_* should comprise two partition coefficients (i.e. core/shell and shell/solvent). Detailed derivations of this mathematical model are given as a **Supplementary Method**. Using MATLAB, the empirical data was fit to Eq. 2 and 3. As shown in **Figure 5B** and **C**, trendline fit for the single-day dataset was better than for the two-day dataset. This was expected, since the daily reservoir change was not accounted for in the two-day case (see **Methods Section** for details). Regardless, the fitted value for diffusion rate constant, *k*_*D*_, was similar for both cases and averaged at 0.04 h^−1^. The best fit *K*_*p*_ for the single-day dataset (31.7) differed from that of the two-day dataset (66.5), though the mean still indicated strong (~50x) DOX preference for 3HM over bulk solvent. Simulations for other *K*_*p*_ values are included for comparison. The expected curve for free DOX (*K*_*p*_= 1) indicated that only <25% DOX content would remain by 36 h. In all three cases, predicted DOX concentration in the reservoir have not yet reached saturation (**Figure 5D**).

Based on scattering results shown earlier, minimal DOX resides in the micelle shell, *C*_*s*_^*m*^vs. the core, *C*_*c*_^*m*^. Hence, partition coefficients for DOX in 3HM core/shell (K_p1_ = *C*_*c*_^*m*^/*C*_*s*_^*m*^) and 3HM shell/bulk solvent or buffer (K_p2_ = *C*_*s*_^*m*^/*C*_*w*_) must be determined (**Figure 5F**). To delineate intra-micellar DOX concentrations, 3HM-DOX samples were repeatedly spin-filtered to remove loosely-bound DOX. Peptide (A280) and DOX (A480) UV-vis absorbance was tracked throughout the spin filtration (**Figure 5E**). After eight spins on Day 1, the A480/280 ratio decreased to a plateau. The sample was stored overnight at 5°C, and spins were resumed the next day. Interestingly, A480/280 ratio dropped further to a second plateau.

The first A480/280 plateau likely indicated complete removal of loosely-bound DOX from the shell (**Figure 5E scheme i - ii**). DOX re-partitioning from the core to shell during storage would enable the second drop and plateau, when more DOX was removed by subsequent spins (**Figure 5E scheme iii - v**). The difference in DOX loading between spin #6 and #16 (0.3%) thus gave the shell-partitioned DOX amount, and loading at spin #16 gave the amount remaining in the core (7.1%). These values yielded K_p1_ = 7.1/0.3, indicating ~ 24x preference of DOX for 3HM core vs. shell. From the dialysis experiment, the overall K_p_ was estimated to be ~50 (comprised of (core: 24 + shell: 1) / solvent = 25/0.5). Hence, K_p2_ ≈ 1/0.5, indicating DOX has a two-fold preference for 3HM shell over bulk solvent. The overall 3HM-DOX partitioning behavior is summarized in **Figure 5F**.

DSC thermograms of samples post-spin #3, 6, and 16 (**Supplementary Figure S4**) deconvoluted to multiple peaks that show successively lower T_m_ approaching that of unloaded 3HM. This indicates sub-populations with decreasing states of alkyl tail crystallinity, as DOX partitioned away from the 3HM core and the loosely-bound molecules were removed at successive spins.

Having a large partition coefficient between DOX in 3HM over bulk solvent is consistent with the slow-release behavior of 3HM-DOX. As shown earlier, DOX release was minimal in non-sink conditions at 35°C (**Figure 3**). Prior studies of 3HM-DOX showed similar slow-release profiles in BSA-sink conditions (~15% release in 20 h^5^), and that 3HM(C16) releases DOX faster than 3HM(C18) during dialysis. ^36^ Repeat FL release experiment with BSA showed ~20% DOX release in 120 h, with t_1/2_ = 36.5 h **(Supplementary Figure S5**), confirming slow DOX release was sustained over extended periods.

DOX’s preferential bias for the 3HM core supports the theory that significant core disruption is necessary for promoting “burst release”, owing to planar DOX intercalation among crystallized alkyl chains. Accordingly, no appreciable 3HM-DOX FL release was observed until solution temperature exceeded the alkyl tails melting temperature of ~40°C (see **Figure 3** and **ref. 5**). Alternatively, “burst release” can be achieved when the amphiphile peptide headgroups are cleaved by proteases (e.g. in lysosomes), causing the whole micelle to lose structural integrity.^5^

Having demonstrated the interplay between DOX location, intra-micellar partitioning, co-assembly energies, and release kinetics, it is evident that 3HM-DOX possesses a harmonized suite of physicochemical characteristics. Detailing the structural, kinetic and thermodynamic nature of cargo-carrier assembly will aid in interpreting behavior in more complex environments. *In vivo*, where physiological temperatures ~37°C remain lower than the alkyl tail melting temperature, strong interactions between DOX molecules and the 3HM core is expected to hold true. In combination with tumor transport parameters, ^72^ cell internalization kinetics, and drug activity time-course, ^73^ ongoing efforts are being pursued to project 3HM-DOX release post-injection and estimate time-cytotoxicity curves in tumor cells. In this way, recommendations for optimal dosing strategies can be made, which maximizes anti-tumor activity while minimizing side effects. Studying nanocarriers with molecular precision can also help to determine its merits as a therapeutic candidate, circumventing wasted resources in preclinical and clinical studies. Hopefully, other drug nanocarrier researchers can appreciate the level of detail and experimental approaches demonstrated here as requisite for sufficient system characterization.

## 4. Conclusions

In summary, fundamental knowledge has been developed on a sub-20 nm nanocarrier-cargo interactions, dimensions, and energy. Present results have answered the “where”, “how much”, and “when” cargo loading and release occurs, which is strongly tied to geometric and energetic considerations of the co-assembly. DOX was found to intercalate into the 3HM alkyl-tail core close to the core-shell interface, maintaining micelle size and uniformity, due to its favorable geometry and hydrophobicity index (GPP*Log P < 1.5). This intercalation promoted higher ordering of the crystalline alkyl tails (T_m_ increased by 8°C), and resulted in a 62% increase of micelle dissociation E_a_ compared to the unloaded 3HM (193 to 312 kJ/mol). In formulation solutions, appreciable DOX release only occurred at T ≥ 45°C with an E_a_ of 83 kJ/ mol. This high energy barrier explains the slow cargo release behavior of 3HM-DOX in physiological environments, with t_1/2_ ~ 40 h in either dialysis reservoir or BSA sink conditions. DOX partitions with ~50x preference for 3HM over the solvent, with K_p1_ core/shell ~24 and K_p2_ shell/solvent ~2. 3HM-DOX’s stability and slow release explains previously-observed long *in vivo* circulation, which is advantageous for ongoing preclinical studies. Molecular-level understanding of the interplay between micellar internal structure, drug co-assembly, release kinetics and thermodynamics have created a detailed picture of the nanocarrier platform and tunable design aspects. These in-depth insights into nanoparticle-cargo co-assembly will lead to eventual predictive design and engineering controls, applicable to the broader nanomedicine drug development process.

## ASSOCIATED CONTENT

### Supplementary Material

Supplementary Material to this article is available online at DOI: j.jconrel. XXXXXXX.

## AUTHOR INFORMATION

### Corresponding Author

* Mailing Address: 381 Hearst Memorial Mining Building, UC Berkeley, Berkeley, CA 94720.

### Author Contributions

B.T.J., M. Lim and T.X. formulated the project. B.T.J., M. Lim, K.J., and M. Li synthesized and characterized materials used. B.T.J. and M. Lim contributed equally to this work. B.T.J. performed kinetic experiments, scattering analysis, and release experiments. M. Lim performed drug property analysis, release experiments / modeling, and partitioning experiments. M. Li developed the MATLAB code for dialysis release and partitioning. N.D. performed loading experiments with small molecules besides DOX. H.D. prepared samples for and executed SANS experiments. T.X. guided project progress. The manuscript was written through contributions of all authors. All authors have given approval to the final version of the manuscript.

### Notes

The authors declare no competing financial interest.

## Acknowledgements

We acknowledge the funding support from Tsinghua-Berkeley Shenzhen Institute (TBSI). B.T.J. was supported by an NSF Graduate Fellowship (DGE 1752814). M. Lim was supported by the UC Berkeley Chancellor’s Fellowship. SAXS data was collected on the X-ray Operations and Research beamline 8-ID-I at the Advanced Photon Source, Argonne National Laboratory. Use of the Advanced Photon Source was supported by the U. S. Department of Energy, Office of Science, Office of Basic Energy Sciences, under Contract No. DE-AC02-06CH11357. We acknowledge the help of Zhiyuan Ruan in SAXS data collection. SANS data was collected at beamline NG3 at the National Institute of Standards and Technology (Gaithersburg, MD) We acknowledge the support of the National Institute of Standards and Technology, U.S. Department of Commerce, in providing the neutron facilities used in this work. This work utilized facilities supported in part by the National Science Foundation under Agreement No. DMR-1508249. We also thank Matthew Bellamy (Univ. of AR Little Rock) for assistance with mathematical model derivations and geometric model approximation.

