## Supplementary Methods and Data for "Designing Sub-20 nm Nanocarriers for Small Molecule Delivery: Interplay among Structural Geometry, Assembly Energetics, and Cargo Release Kinetics"

Electronic Supplementary Material

Supplementary Methods

### 3HM-DOX Dialysis Release Profiles

#### I. Differential Equation Derivations for One Partition Coefficient

Consider a case where DOX partitions between the micelle and bulk solvent (water) inside the dialysis chamber (Figure 1). The only driving force for DOX removal is diffusion across the dialysis membrane, moving free DOX from the water to the reservoir.

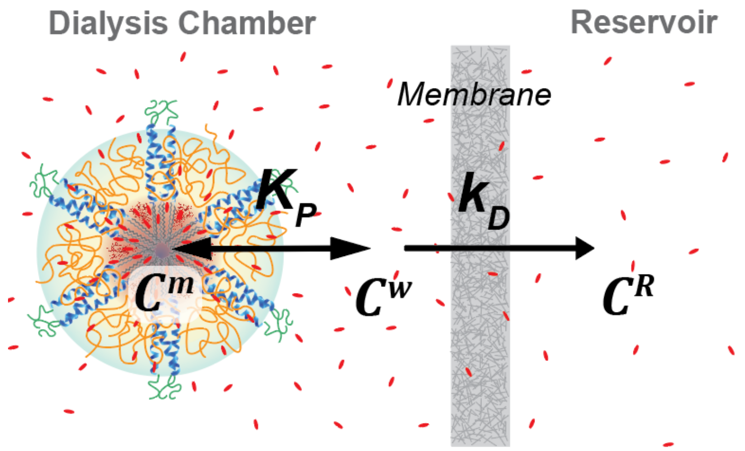


Figure 1: Schematic of DOX release with one partition coefficient

The mass balance of total DOX ($M^T$) inside dialysis chamber, accounting for DOX inside micelle ($M^m$), and bulk solvent ($M^w$), is given by:

$$M^T = M^m + M^w \tag{1}$$

We divide by the total chamber volume ($V^T$) to switch from total DOX mass to concentration ($C^T$), and convert the other mass terms to concentrations ($C^m$, $C^w$) by applying volume fractions for micelle ($V^m$) and bulk solvent ($V^w$):

$$C^T = \frac{M^T}{V^T} = \frac{V^m}{V^T} C^m + \frac{V^w}{V^T} C^w \tag{2}$$

The DOX partition coefficient between micelle and bulk solvent is given by:

$$K_p = C^m / C^w \tag{3}$$

The diffusion rate equation for 3HM-DOX dialysis is described using rate constant $k_D$. The only driving force is free DOX moving out of the bulk solvent into reservoir (Figure 1). The following is thus the rate of change of DOX concentration in the chamber:

$$\frac{\mathrm{d}C^T}{\mathrm{d}t} = -k_D \left( C^w - C^R \right) \tag{4}$$

Similarly, we set up a rate of change equation for DOX concentration in the reservoir, accounting for the volume ratio between the dialysis chamber and reservoir:

$$\frac{\mathrm{d}C^R}{\mathrm{d}t} = \frac{V^T}{V^R} k_D \left( C^w - C^R \right) \tag{5}$$

Ultimately, we only measured $C^T$ over time, hence we need Equations (5) and (6) to be written in terms of $C^T$. To do so, we must consolidate all the concentration terms by incorporating the partition coefficient.

Rearrange Equation (3) to obtain a single expression for $C^w$ in terms of $C^m$ using the partition coefficient:

$$C^w = \frac{C^m}{K_p} \tag{6}$$

Substitute Equation (6) to the rate Equations (4) and (5):

$$\frac{\mathrm{d}C^T}{\mathrm{d}t} = -k_D \left( \frac{C^m}{K_p} - C^R \right) \tag{7}$$

$$\frac{\mathrm{d}C^R}{\mathrm{d}t} = \frac{V^T}{V^R} k_D \left( \frac{C^m}{K_p} - C^R \right) \tag{8}$$

Then, we also substitute Equation (6) to the original mass balance Equation (2), to relate $C^m$ in terms of $C^T$:

$$C^T = \frac{V^m}{V^T} C^m + \frac{V^w}{V^T} \frac{C^m}{K_p} \tag{9}$$

$$C^T = C^m \left( \frac{V^m}{V^T} + \frac{V^w}{V^T} \frac{1}{K_p} \right) \tag{10}$$

$$C^m = C^T / \left( \frac{V^m}{V^T} + \frac{V^w}{V^T} \frac{1}{K_p} \right) \tag{11}$$

Finally, we substitute Equation (11) into rate Equations (7) and (8):

$$\frac{\mathrm{d}C^T}{\mathrm{d}t} = -k_D \left( \frac{C^T}{K_p \frac{V^m}{V^T} + \frac{V^w}{V^T}} - C^R \right) \tag{12}$$

$$\frac{\mathrm{d}C^R}{\mathrm{d}t} = \frac{V^T}{V^R} k_D \left( \frac{C^T}{K_p \frac{V^m}{V^T} + \frac{V^w}{V^T}} - C^R \right) \tag{13}$$

Simplifying volume fractions as $a = V^m/V^T$ and $b = V^w/V^T$, we arrive at the final equations for the model:

$$\frac{\mathrm{d}C^T}{\mathrm{d}t} = -k_D \left( \frac{C^T}{aK_p + b} - C^R \right) \tag{14}$$

$$\frac{\mathrm{d}C^R}{\mathrm{d}t} = \frac{V^T}{V^R} k_D \left( \frac{C^T}{aK_p + b} - C^R \right) \tag{15}$$

#### Boundary Conditions

B.C. 1: @ t=0, $C^R = 0$

B.C. 2: @ t=0, $C^T = C_0^T$ (initial DOX concentration in sample)

#### Constants

Values for $V^T$, and $V^R$ are determined by the experimental setup.

Volume fractions in $a$ and $b$ are computed using micellar dimensions and concentration (i.e. micelle volume $V_{3HM}$ for 18 nm 3HM at concentration $C_0^T$ mg/mL). $C_0^T$ must be converted to a molar basis before computing the number of micelles per volume via Avogadro's number ($N_{av}$). 3HM molecular weight ($W_{3HM}$) is based on the aggregation number and monomer MW: $45 * 7300\ Da = 328,500\ Da$ (*ref: Dong et al, JACS 2012, 134(28): 11807-11814*).

$$V^m = \frac{C_0^T}{W_{3HM}} N_{av} V_{3HM} V^T \tag{16}$$

$$V^w = V^T - V^m \tag{17}$$

The system of differential equations (14) and (15) will be solved for $k_D$ and $K_p$.

#### II. Differential Equation Derivations for Two Partition Coefficients

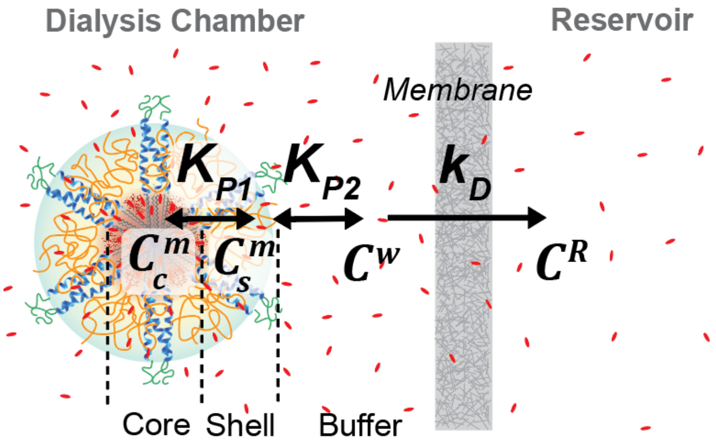


Figure 2: Schematic of DOX release with 2 partition coefficients

Consider a case where DOX partitions between the micelle core and shell, and between the shell to the buffered bulk solvent (water) inside the dialysis chamber (Figure 2). The only driving force for DOX removal is diffusive flux acting across the dialysis membrane, moving free DOX from the water to the reservoir.

Mass balance of total DOX ($M^T$) inside dialysis chamber, accounting for DOX inside micelle core ($M_c^m$), shell ($M_s^m$), and bulk solvent ($M^w$):

$$M^T = M_c^m + M_s^m + M^w \tag{18}$$

Divide by total chamber volume ($V^T$) to switch from DOX total mass to concentration ($C^T$), and convert the other mass terms to concentrations ($C_c^m$, $C_s^m$, $C^w$) by applying volume fractions for micelle core ($V_c^m$), shell ($V_s^m$), and bulk solvent ($V^w$):

$$C^T = \frac{M^T}{V^T} = \frac{V_c^m}{V^T} C_c^m + \frac{V_s^m}{V^T} C_s^m + \frac{V^w}{V^T} C^w \tag{19}$$

Partition coefficient of DOX in core to shell:

$$K_{p1} = C_c^m / C_s^m \tag{20}$$

Partition coefficient of DOX in shell to water:

$$K_{p2} = C_s^m / C^w \tag{21}$$

Rate equation for DOX leaving the sample. The only driving force is free DOX moving out of water into reservoir. The following is thus the rate of change of DOX concentration in the water:

$$\frac{\mathrm{d}C^T}{\mathrm{d}t} = -k_D \left(C^w - C^R\right) \tag{22}$$

Similarly, use set up a rate of change equation for DOX concentration in the reservoir, accounting for the volume ratio between the dialysis chamber and reservoir:

$$\frac{\mathrm{d}C^R}{\mathrm{d}t} = \frac{V^T}{V^R} k_D \left(C^w - C^R\right) \tag{23}$$

Ultimately we only measured $C^T$ over time, hence we need Equations (22) and (23) to be written in terms of $C^T$.

Rearrange partition coefficient Equations (20) and (21) to obtain a single expression for $C^w$:

$$C_s^m = \frac{C_c^m}{K_{p1}} \tag{24}$$

$$K_{p2} = \frac{C_c^m / K_{p1}}{C^w}$$

$$C^w = \frac{C_c^m}{K_{p1} K_{p2}} \tag{25}$$

As shown in Equation (25), we can obtain an expression for $C^w$ in terms of $C_c^m$. Thus, we first substitute Equation (25) to the rate Equations (22) and (23):

$$\frac{\mathrm{d}C^T}{\mathrm{d}t} = -k_D \left( \frac{C_c^m}{K_{p1} K_{p2}} - C^R \right) \tag{26}$$

$$\frac{\mathrm{d}C^R}{\mathrm{d}t} = \frac{V^T}{V^R} k_D \left( \frac{C_c^m}{K_{p1} K_{p2}} - C^R \right) \tag{27}$$

We will also substitute Equation (25) to the original mass balance Equation (18), to relate $C^T$ in terms of $C_c^m$:

$$C^T = \frac{V_c^m}{V^T} C_c^m + \frac{V_s^m}{V^T} \frac{C_c^m}{K_{p1}} + \frac{V^w}{V^T} \frac{C_c^m}{K_{p1} K_{p2}} \tag{28}$$

$$C^T = C_c^m \left( \frac{V_c^m}{V^T} + \frac{V_s^m}{V^T} \frac{1}{K_{p1}} + \frac{V^w}{V^T} \frac{1}{K_{p1} K_{p2}} \right) \tag{29}$$

Simplify with a variable substitution, $\beta$:

$$\beta = \frac{V_c^m}{V^T} + \frac{V_s^m}{V^T} \frac{1}{K_{p1}} + \frac{V^w}{V^T} \frac{1}{K_{p1} K_{p2}} \tag{30}$$

Hence:

$$C_c^m = C^T / \beta \tag{31}$$

Finally, we substitute Equation (31) into Equations (26) and (27). These are the final equations for the model:

$$\frac{\mathrm{d}C^T}{\mathrm{d}t} = -k_D \left( \frac{C^T / \beta}{K_{p1} K_{p2}} - C^R \right) \tag{32}$$

$$\frac{\mathrm{d}C^R}{\mathrm{d}t} = \frac{V^T}{V^R} k_D \left( \frac{C^T/\beta}{K_{p1} K_{p2}} - C^R \right) \tag{33}$$

#### Boundary Conditions

B.C. 1: @ t=0, $C^R = 0$

B.C. 2: @ t=0, $C^T = C_0^T$ (initial DOX concentration in sample)

#### Constants

Values for $V^T$ and $V^R$ are determined by the experimental setup.

Volume fractions in $\beta$ are computed using micellar dimensions and concentration (i.e. micelle volume $V_{3HM}$ for 18 nm 3HM, with core and shell radii given in the main text for $V_{core}$ and $V_{shell}$, at concentration $C_0^T$ mg/mL). $C_0^T$ must be converted to a molar basis before computing the number of micelles per volume via Avogadro's number $N_{av}$. 3HM molecular weight ($W_{3HM}$) is based on the aggregation number and monomer MW: $45 * 7300\ Da = 328,500\ Da$ (*ref: Dong et al, JACS 2012, 134(28): 11807-11814*).

$$V_c^m = \frac{C_0^T}{W_{3HM}} N_{av} V_{core} V^T \tag{34}$$

$$V_s^m = \frac{C_0^T}{W_{3HM}} N_{av} V_{shell} V^T \tag{35}$$

$$V^w = V^T - V_c^m - V_s^m \tag{36}$$

The system of differential equations (32) and (33) can be solved for $k_D$, $K_{p1}$ and $K_{p2}$.

#### Supplementary Tables

**Table S1**: Calculated X-ray SLDs of 3HM-DOX components.

| X-Ray | SLD ($10^{-6}$ Å$^{-2}$) | Density (g/cm$^3$) |
|---|---|---|
| 1CW peptide | 13.2 | 1.40 |
| C18 alkyl | 9.33 | 0.94 |
| PEG | 11.5 | 1.20 |
| DOX | 14.9 | 1.60 |
| $H_2O$ buffer | 9.52 | 1.00 |

**Table S2**: Calculated neutron SLDs of 3HM-DOX components. SLDs are given for hydrogenated (H) vs. deuterated (D) alkyl tails.

| Neutron | SLD ($10^{-6}$ Å$^{-2}$) | Density (g/cm$^3$) |
|---|---|---|
| 1CW peptide | 1.77 | 1.40 |
| C16 alkyl (H/D) | -0.08/6.63 | 0.85 /0.96 |
| PEG | 0.67 | 1.20 |
| DOX | 2.56 | 1.60 |
| $D_2O$ buffer | 6.39 | 1.11 |

**Table S3**: Fitted X-ray SLDs of 3HM-DOX core and shell.

| Sample | 3HM (C18) | 3HM-DOX 6 % | 3HM-DOX 7.2 % | 3HM-DOX 8 % |
|---|---|---|---|---|
| Core SLD ($10^{-6}$ A$^{-2}$ ) | 10.5 | 12.8 | 13.6 | 13.7 |
| Shell SLD ($10^{-6}$ A$^{-2}$ ) | 9.9 | 9.9 | 10.0 | 10.1 |

**Table S4**: Fitted neutron SLDs of 3HM-DOX core and shell. SLDs are given for hydrogenated (H) vs. deuterated (D) alkyl tails.

| Sample | 3HM (C16) | 3HM-DOX (C16) |
|---|---|---|
| Core SLD (H/D) ($10^{-6}$ A$^{-2}$ ) | -0.08/6.0 | 0.68/5.0 |
| Shell SLD (H/D) ($10^{-6}$ A$^{-2}$ ) | 5.4/5.9 | 4.9/5.2 |

**Table S5:** Rate constants calculated from fitting data shown in **Figure 4B** to facilitate the calculation of kinetic $E_a$.

| Temp (°C) | Inflection Point (°C) | Rate, k ($h^{-1}$) | $t_{1/2}$ (h) |
|---|---|---|---|
| 55 | 16.9 | .0021 | 330 |
| 60 | 15.2 | .0051 | 136 |
| 65 | 10.1 | .0098 | 71 |
| 70 | 7.9 | .015 | 46 |

**Table S6:** Phase transition temperatures and total enthalpies of 3HM-DOX over a range of scan rates corresponding to data shown in **Figure 4D**.

| Scanrate (ºC/min) | $T_m$ (°C) | ΔH (kJ/mol) |
|---|---|---|
| 0.1 | 30.6 | 29.8 |
| 0.5 | 35.5 | 34.9 |
| 0.75 | 36.2 | 36.3 |
| 1 | 36.3 | 34.6 |
| 1.25 | 36.5 | 33.8 |
| 1.5 | 37.0 | 36.0 |

#### Supplementary Figures

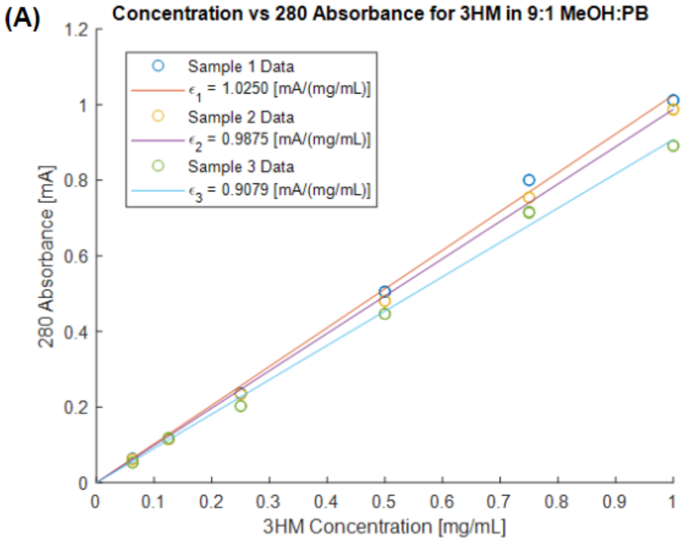


| Sample | $\varepsilon_{280}$ [cm$^{-1}$mg/mL$^{-1}$] |
|---|---|
| 1 | 1.025 |
| 2 | 0.988 |
| 3 | 0.908 |
| Average | 0.974 |

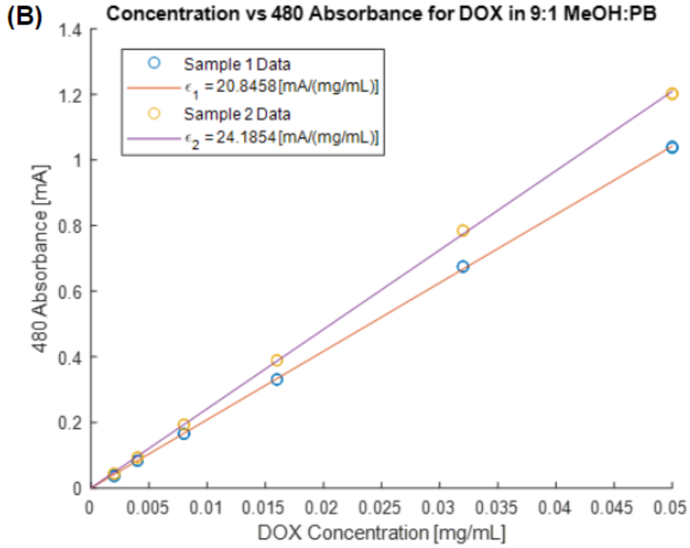


| Sample | $\varepsilon_{480}$ [cm$^{-1}$mg/mL$^{-1}$] |
|---|---|
| 1 | 20.8 |
| 2 | 24.2 |
| Average | 22.5 |

**Figure S1:** Extinction coefficient curves for (A) 3HM A280 and (B) DOX A480.

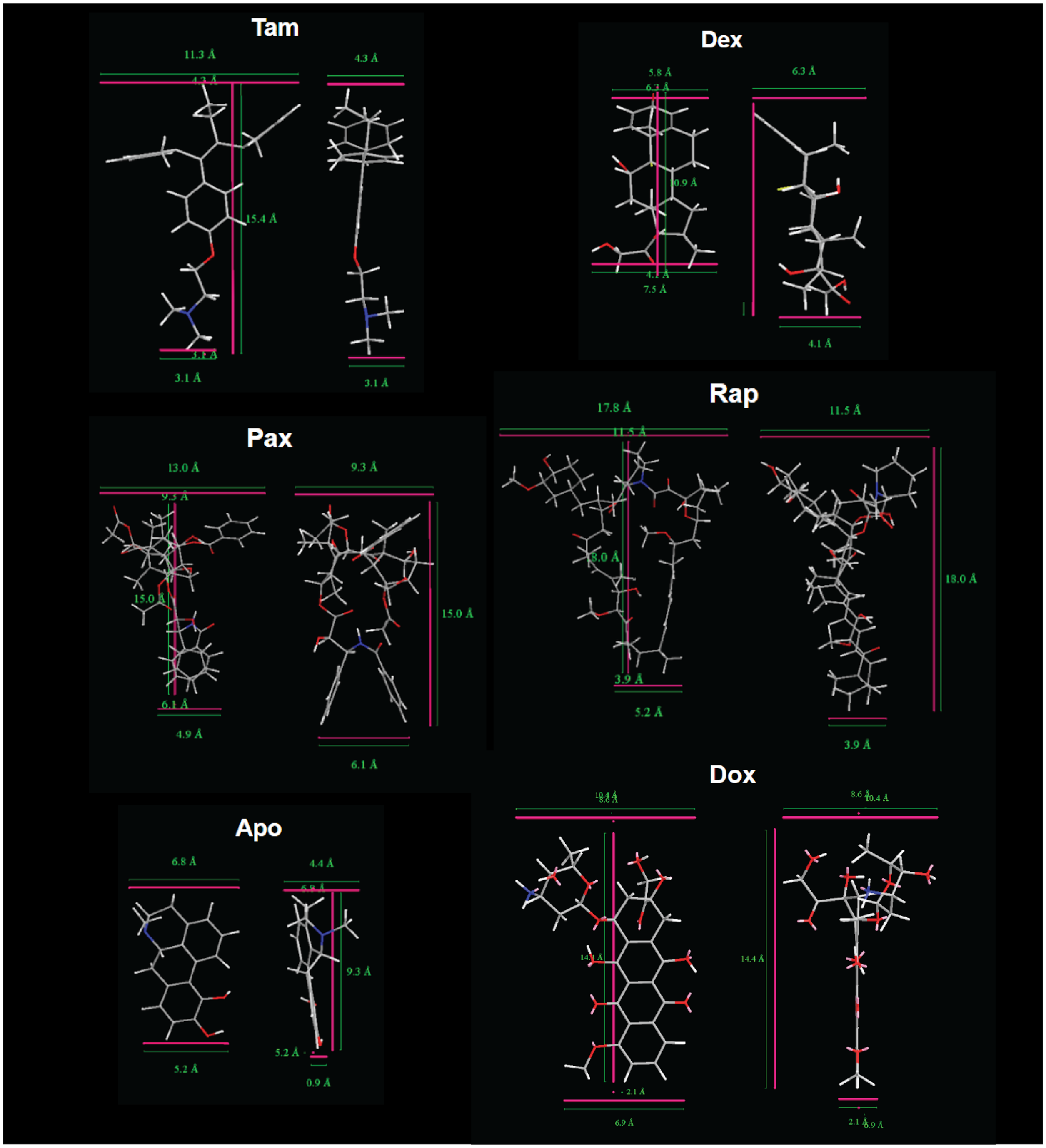


**Figure S2:** Small molecule drug Chem3D structure measurements.

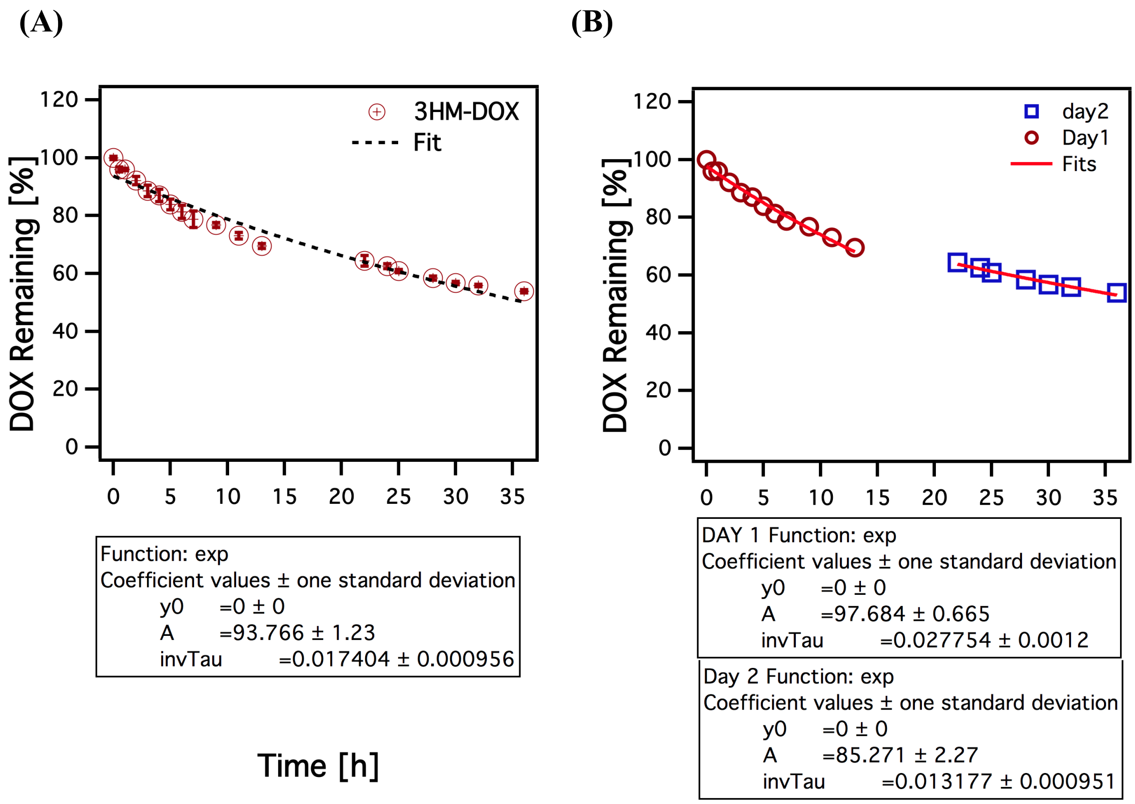


**Figure S3**: Dialysis release fits with negative exponential function (A) over the entire time duration ($t_{1/2}$ = 39.8 h), and (B) for each of the two-day experiment (Day 1 release $t_{1/2}$ = 25.0 h, Day 2 $t_{1/2}$ = 52.5 h, averaging 39.8 h).

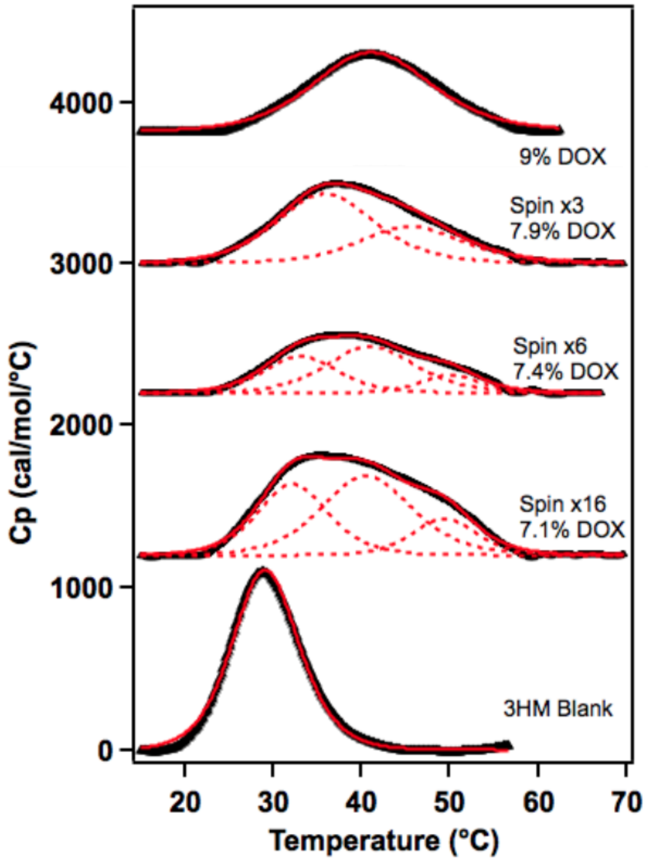


**Figure S4**: DSC of 3HM-DOX samples after successive deionized water washes via spin filtration.

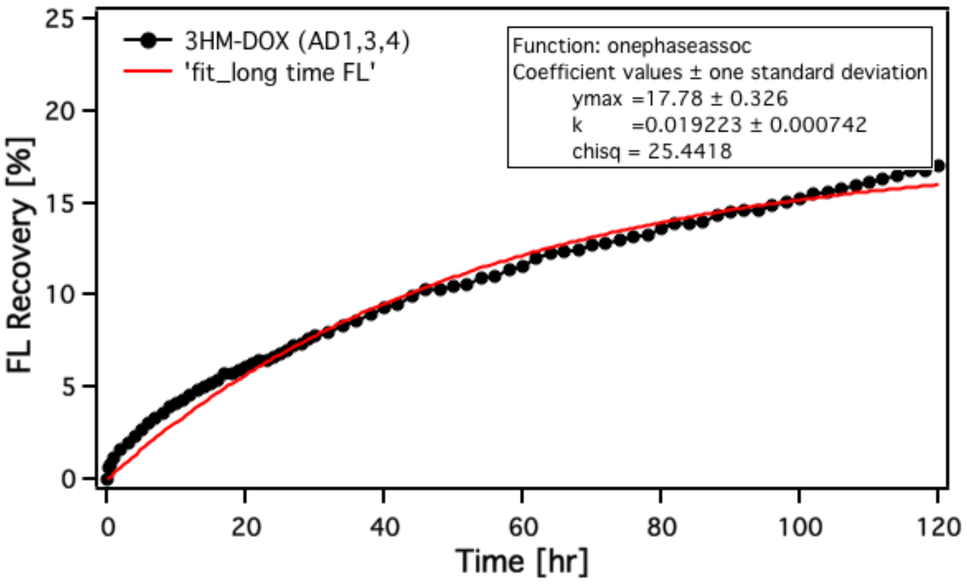


**Figure S5:** FL release of DOX from 3HM-DOX in buffer containing 50 mg/ mL BSA at 37 °C.
